# Thermodynamics and kinetics of the amyloid-β peptide revealed by Markov state models based on MD data in agreement with experiment

**DOI:** 10.1101/2020.07.27.223487

**Authors:** Arghadwip Paul, Suman Samantray, Marco Anteghini, Birgit Strodel

**Author notes:** Institute of Biological Information Processing: Structural Biochemistry (IBI-7), Forschungszentrum Jülich, 52428 Jülich, Germany. Institute of Theoretical and Computational Chemistry, Heinrich Heine University Düsseldorf, 40225 Düsseldorf, Germany (Birgit Strodel). LifeGlimmer GmbH, Markelstraße 38 12163 Berlin, Germany (Marco Anteghini).

## Abstract

The amlyoid-β peptide (Aβ) is closely linked to the development of Alzheimer’s disease. Molecular dynamics (MD) simulations have become an indispensable tool for studying the behavior of this peptide at the (sub)molecular level, thereby providing insight into the molecular basis of Alzheimer’s disease. General key aspects of MD simulations are the force field used for modeling the peptide or protein and its environment, which is important for accurate modeling of the system of interest, and the length of the simulations, which determines whether or not equilibrium is reached. In this study we address these points by analyzing 30-µs MD simulations acquired for Aβ40 using seven different force fields. We assess the convergence of these simulations based on the convergence of various structural properties and of NMR and fluorescence spectroscopic observables. Moreover, we calculate Markov state models for each of the seven MD simulations, which provide an unprecedented view of the thermodynamics and kinetics of the amyloid-β peptide. This further allows us to provide answers for pertinent questions, like: *Which force fields are suitable for modeling Aβ?* (a99SB-UCB and a99SB-ILDN/TIP4P-D); *What does Aβ peptide really look like?* (mostly extended and disordered) and; *How long does it take MD simulations of Aβ to attain equilibrium?* (20–30 µs). We believe the analyses presented in this study will provide a useful reference guide for important questions relating to the structure and dynamics of Aβin particular, and by extension other similar disordered peptides.

## 1 Introduction

The amyloid-β peptide (Aβ) plays a central role in the development of Alzheimer’s disease (AD), a neurodegenerative disease that currently affects about 50 million people worldwide.^1^ This number is expected to almost double every 20 years, reaching 75 million in 2030 and 131.5 million in 2050.^1^ Despite intense search for a cure for AD, there is currently no targeted therapy able to stop or slow its progression; only drugs that may help treat symptoms for a couple of months are available. The development of AD drugs is characterized by an exceptionally high rate of failure. Between 2002 and 2012, 99.6% of drug trials aimed at preventing, curing or improving the symptoms of AD had failed or been discontinued.^2^ To put this into context, the failure rate for cancer drugs is 81%. ^3^ Lessons learned from several years of research into AD treatment shows that a good understanding of the structural and molecular basis of AD pathophysiologic mechanisms is a prerequisite for the rational design of AD drugs.

The aggregation of Aβ is closely linked to the onset and development of AD;^4^ it is thus of paramount importance to unravel the details of this process. In the past two decades a plethora of biochemical and biophysical studies have been conducted that were aimed at exactly that goal.^5^ Nonetheless, for various reasons many open questions surrounding the aggregation of Aβ remain. The difficulties in characterizing this process lie in the facts that Aβ is a highly flexible peptide belonging to the class of intrinsically disordered proteins (IDPs) and can follow different aggregation pathways, which may be on- or off-pathway towards the end products of the amyloid aggregation route that are amyloid fibrils composed predominantly of β-sheet structure in a characteristic cross-β conformation. Moreover, different amyloid fibril structures exist for Aβ and an even higher level of polymorphism is found for the intermediate aggregation products. Other complicating aspects are that the aggregation process operates on a huge range of length- and timescales and is highly sensitive to external conditions.^6^ Moreover, the aggregation behavior depends on the particular alloform of Aβ, which exists in different lengths ranging from of 36 to 43 amino acids that are found as the main components of the amyloid plaques populating the brains of people having AD.^7^ The two major alloforms are those with 40 and 42 residues, denoted as Aβ40 and Aβ42 in the following. The most abundant form is Aβ40, while Aβ42 is more prone to aggregate and therefore more frequently deposited in amyloid plaques.^7^

Among the multitude of physicochemical techniques that are employed for studying Aβ, molecular dynamics (MD) simulations at the atomic level provide the highest spatial and temporal resolution for capturing the structural and dynamical characteristics of this peptide.^8^ Many of the simulation studies, yet as also many of the experimental investigations of Aβ are focused on its monomeric state since the properties of the monomers of aggregation-prone peptides and proteins are the determinants of their aggregation behavior.^9,10^ It has been 15 years since the first all-atom MD simulation of full-length Aβ in solution has been performed.^11^ Since then, hundreds of simulation studies involving Aβ40 or Aβ42 have been published. During the first ∼10 years of these studies, research groups relied on the common protein and water force fields (FFs) as was usual practice for the simulation of folded proteins, which were the more frequent object of simulation studies at that time. However, as time showed, very different results regarding the structural preferences of Aβ were obtained depending on which of the FFs was used. In fact, as far back as 2012 when we had first reported our first simulation study of Aβ in solution we concluded that ‘proper benchmarking of the protein force fields for unfolded and intrinsically disordered proteins’ was needed. ^12^ Over the years, various FF benchmarks for Aβ and IDPs in general have been performed.^13–17^ Depending on the FFs tested, the simulation technique employed (i.e., standard MD versus enhanced-sampling MD, like replica exchange MD), the length of the simulations, and the experimental data used for validation, different FFs were identified as the most suited ones. For instance, García et al.^13,14^ found OPLS-AA^18,19^ with the TIP3P water model^20^ and AMBER99SB^21^ with the TIP4P-Ew water model^22^ to be the best FFs for Aβ42, while our own benchmark, that included AMBER99SB,^21^ AMBER99SB*-ILDN, ^23,24^ AMBER99SB-ILDN-NMR,^25^ and CHARMM22*^26^ combined with TIP4P-Ew^22^ as well as OPLS-AA/TIP3P,^18–20^ identified CHARMM22* as the best FF.^17^ However, more recent benchmarks revealed that the common FFs from the Amber, GROMOS, OPLS, or CHARMM family in combination with standard three- or four-point water models produce conformational ensembles for IDPs that are too compact and too biased towards folded structures.^27,28^ This conclusion can be explained with the parametrization strategy underlying the standard FFs, which aimed at producing the correct structure for folded proteins,^29^ while IDPs do not adopt a well-defined equilibrium structure in solution, instead sampling an ensemble of fully and/or partially disordered structures.

A number of research groups set out to modify the existing FF parameters such that they capture the structural diversity and flexibility of IDPs, producing less folded and more expanded IDP conformations. These force field modifications include strengthening the water-protein London dispersion interactions,^28,30^ refining the protein backbone parameters to create more expanded structures or reducing the tendency for certain ordered conformations,^31^ and/or altering the salt-bridge interactions.^31^ In a recent effort, D. E. Shaw Research used AMBER99SB*-ILDN^23,24^ combined with TIP4P-D water^28^ as a starting point and reparametrized torsion parameters and the protein–water van der Waals (vdW) interaction terms with the aim to develop a FF that provides an accurate model for both folded proteins and IDPs.^32^ The performance of the resulting FF, called a99SB-disp, was tested for a large benchmark set of 21 experimentally well-characterized proteins and peptides, including folded proteins, fast-folding proteins, weakly structured peptides, disordered proteins with some residual secondary structure, and disordered proteins with almost no detectable secondary structure. In addition, they assessed the accuracy of six other FF/water combinations. Two of these combinations belong to the older FFs that were developed for folded proteins, a99SB*-ILDN/TIP3P (‘a’ standing for AMBER)^23,24^ and C22*/TIP3P (‘C’ for CHARMM)^26^ and three combinations specifically designed for IDPs: a03ws containing empirically optimized solute–solvent dispersion interactions,^30^ a99SB-ILDN/TIP4P-D with increased water dispersion interactions,^28^ and C36m with refined backbone potentials (which by default is used with CHARMM-modified TIP3P).^31^ The seventh FF in the benchmark is a99SB-UCB, which is based on a99SB/TIP4P-Ew^21,22^ and includes modified backbone torsion parameters^33^ and optimized protein–solvent Lennard-Jones (LJ) parameters^34^ proposed by Head-Gordon and co-workers. For each of the proteins or peptides in their test set and FFs considered, Robustelli et al. performed 30 µs MD simulations and compared the MD-generated ensembles against a number of experimental data mainly derived from nuclear magnetic resonance (NMR) sprectroscopy, and, if available, also from small angle X-ray scattering (SAXS) and fluorescence resonance energy transfer (FRET).^32^ They concluded that, taking all tested proteins and peptides into consideration, a99SB-disp is the best-performing FF. One of the peptides included in their test set is Aβ40. Based on Figure 2 of ref 32, a99SB-disp is, together with C22*/TIP3P, the second best choice following C36m for modeling Aβ40. Interestingly, the performance of a99SB-UCB is not shown in this figure. Though for Aβ40 it was concluded that ‘a99SB-UCB produced excellent agreement with experimental NMR measurements’.^32^

**Figure 1:**
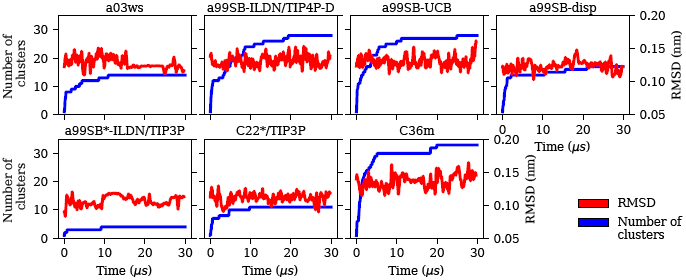
Evolution of the C_α_-RMSD with respect to the Aβ40 starting structure of the MD simulations (red, right *y*-axis) and the number of conformational clusters (blue, left *y*-axis) for the different force fields (labels on the top of the panels).

**Figure 2:**
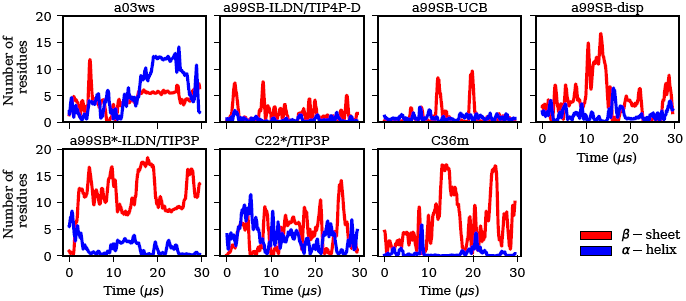
Evolution of the secondary structures β-sheet (red) and α-helix (blue) in terms of the number of Aβ40 residues adopting these structures for the different force fields (labels on the top of the panels).

A question that was not addressed by the study of Robustelli et al. is how good the different FFs are able to reproduce the dynamics of the different proteins and peptides. From a FRET study of Aβ40 and Aβ42 it was found that both peptides do not exhibit conformational dynamics exceeding 1 µs,^35^ which also agrees with the findings from fluorescence measurements using the method of Trp-Cys contact quenching. ^36^ Given the simulation length of 30 µs of the MD data generated by D. E. Shaw Research, we use their data (which were kindly provided by them) to assess the kinetics of Aβ40 as sampled by the different FFs. Our primary goal is to determine how much MD sampling is needed to reach convergence with standard MD simulations applied to Aβ. To this end, we evaluate the convergence of intrinsic structural quantities as well as of NMR observables calculated from the MD data. Moreover, we generate Markov state models (MSMs), which, in addition to providing stringent convergence checks, also elucidate the kinetically stable states of Aβ and the transitions between them. This analysis reveals that the length of an MD simulation required for obtaining equilibrated results for Aβ strongly depends on the FF used for modeling the peptide, but usually requires 20 µs or more. Another finding is that only two of the seven FFs under consideration are able to reproduce both the structural and kinetic data available from experiments of Aβ40, which are a99SB-UCB, which performs by far the best, and a99SB-ILDN/TIP4P-D. After many years of research, with a99SB-UCB and tens of microsecond of MD simulations it is now possible to predict the thermodynamics and kinetics of the amyloid-β peptide.

## 2 Methodology

### MD trajectories

The MD trajectories were generated by Robustelli et al.^32^ and kindly provided by the authors. All MD simulations were initiated from an Aβ40 structure similar to that found in PDB entry 1BA6^37^ and run for 30 µs using following FFs: a03ws, a99SB-ILDN/TIP4P-D, a99SB-UCB, a99SB-disp, C22*/TIP3P, and C36m. The following analyses was applied to each of these seven trajectories.

### Structural analyses

The trajectories were analyzed using a combination of standard Gromacs-2016.4 package tools, ^38,39^ custom written Tcl scripts in VMD,^40^ and Python scripts using the MDAnalysis^41^ and MDTraj^42^ libraries. As the trajectory files from the Desmond MD package ^43^ are in DCD file format, there was a need for conversion into Gromacs-compatible TRR format.

After this, protein conformations were clustered using the clustering algorithm of Daura et al.^44^ as implemented in Gromacs with a root mean square deviation (RMSD) cutoff of *R*_cut_ = 1.0 nm. To assign secondary structure elements to the protein conformations, the STRIDE algorithm^45^ was employed. Inter-residue contact maps were constructed by calculating the fraction of structures in which the residue pairs were having at least one pair of atoms within 0.4 nm of each other.

### Construction of Markov state models

The underlying kinetics of the systems are captured by constructing Markov state models (MSMs) from the MD simulation data using the PyEMMA library in Python.^46^ The first step towards building an MSM is to choose a suitable distance metric, called feature, for defining the conformational space of the molecule. Here, we describe the conformations in terms of the distances between the C_*α*_ atoms. This feature was selected based on VAMP-2 scores, where VAMP stands for Variational Approach for Markov Process.^47^ This score is part of the VAMP scores family, which represents a set of score functions that can be used to find optimal feature mappings and optimal Markovian models of the dynamics from time series data. In order to choose a subset of relevant features for our model construction, we considered three different features: C_*α*_ distances, minimum distance between residues, and backbone torsion angles. In order to evaluate which feature is the best and to avoid overfitting, a cross-validation was performed, comparing the VAMP-2 scores of each of the three features computed for three subtrajectories of 10 µs length per force field. From this analysis, C_*α*_ distances emerged as the most suitable feature (data not shown).

Next, we reduced the dimension of the space from 703 interatomic distances to 2 collective coordinates by applying time-lagged independent component analysis (TICA), a dimensionality reduction technique that identifies the slowest modes in the feature space by maximizing the autocorrelation of the reduced coordinates,^48^ and hence is preferred for MSM construction over the more commonly used principal component analysis (PCA), which does not take into account any kinetic information. A density based clustering technique, HDBSCAN^49^ is then applied to decompose the reduced conformation space into a set of disjoint discrete states and define the trajectory as a sequence of transitions between these states. An MSM can next be built from this discrete trajectory by counting the transitions between the states at a specified lag-time, constructing a matrix of the transition counts and normalizing it by the total number of transitions emanating from each state to obtain the transition probability matrix. However, the resulting MSM is too granular to provide a simple, intuitive picture of the system dynamics. That is achieved by coarse-graining the MSM into a hidden Markov model (HMM) with a few metastable states, using robust Perron cluster analysis (PCCA+),^50^ which is a fuzzy version of the spectral algorithm for partitioning graphs that assigns each microstate a probability of belonging to a metastable macrostate. ^51^ Finally, whether the final model satisfies the Markovian assumptions is verified with a Chapman-Kolmogorov test.^52^

In the following, only HMMs are presented, which are, however, for simplicity denoted as MSMs. Thus, all MSM states discussed in this work are coarse-grained states that resulted from PCCA+ applied to the fine-grained MSMs.

### Calculation of experimental observables

The NMR chemical shifts of the protein backbone atoms were calculated using Sparta+^53^. Dihedral angles (ϕ, ψ) were calculated for each Aβ residue from the MD trajectories and converted into residue-specific backbone scalar ^3^*J*_HNH*α*_ coupling constants using the Karplus equation^54^ with the Karplus coefficients values *A* = 7.97 Hz, *B* = −1.26 Hz, and *C* = 0.63 Hz from Vogeli *et. al*.^55^ The simulated and experimentally derived coupling constants were compared by calculating *χ*^2^ using eq (1), including the error term Δ = 0.42 Hz:^30,35^

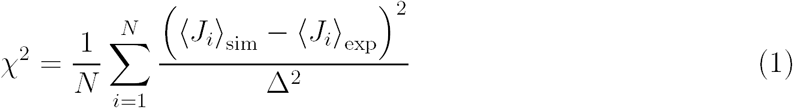

Here, *J*_*i*_ represents the *J* -coupling constant for the *i*-th residue, *N* is the total number of residues for which the experimental data are available, the subscripts “sim” and “exp” correspond to the simulated and experimental data respectively, and *·* denotes the ensemble average.

Time-series of the end-to-end distance *R*_ee_ were calculated as the distance between the *C*_*α*_ atoms of the N- and C-termini of Aβ40 using the MDTraj library^42^ in Python. From this, the FRET efficiency is calculated as

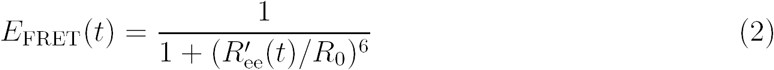

where the Förster radius *R*_0_ = 5.2 nm for the dye pair of Alexa 488 and 647^35,56^ was used, and 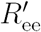 is calculated by scaling up *R*_*ee*_ to account for the effects of the experimental dyes by treating them as 12 extra amino acid residues and assuming a Gaussian scaling exponent,^35^

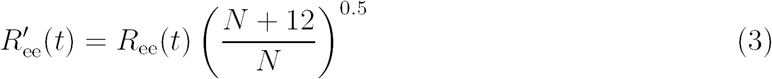

where *N* = 40 is the number of residues in the peptide under study.

### Bayesian reweighting of trajectories by using experimental data

The Bayesian/maximum entropy (BME) technique^57^ was used to reweight the trajectories and obtain a refined conformational ensemble consistent with selected experimental data, thereby compensating for the discrepancies between the experimental and calculated observables which arise from inaccuracies in the force fields. Here, we considered the *J* -coupling data to obtain the optimized set of weights for the a99SB-UCB and C36m trajectories. The regularization parameter *θ* in BME, which balances the trust in data versus simulation, ^58^ was taken as 10.

## 3 Results

We used the 30 µs MD data of Aβ40 from Robustelli et al.^32^ to i) assess the convergence of these trajectories, ii) determine the agreement of the simulated Aβ40 ensembles with spectroscopic data, and iii) derive the thermodynamics and kinetics of this peptide. The convergence was tested for various structural properties that are usually calculated from MD trajectories, including the structural RMSD, clustering analysis, radius of gyration (*R*_gyr_), and the secondary structure (section 3.1). Another kind of convergence check is provided by Markov state models (section 3.2), which is based on kinetically clustering the MD data. Section 3.3 contains the calculation of NMR spectroscopic and FRET observables which allows us to compare the MD generated structural ensembles with experimental findings and to also assess the convergence of these spectroscopic quantities. In section 3.4 we evaluate the kinetics of Aβ40 and compare the MD results to experimental observations. The structural ensemble of Aβ40 in agreement with thermondyanmic and kinetic data derived from experiments is examined in the Discussion following thereafter.

### 3.1 Convergence checks based on structural data

#### RMSD

As commonly done with MD data, we calculated the C_*α*_ RMSD of the 30-µs MD trajectories with respect to the starting structure of these simulations. From the time evolution of the RMSD shown in Figure 1 one can see that the Aβ40 conformations quickly move away from the initial conformation, reaching values of about 1 to 1.5 nm within a few nanoseconds. One can further observe that within the 30-µs of sampling the RMSD does not considerably further increase but fluctuates between 1 and 1.5 nm, or somewhat above 1.5 nm for some of the FFs. Based on the RMSD profiles alone one might easily but incorrectly be tempted to conclude that the simulations converged within a few nanoseconds. As our analyses will show in the following sections, this would have been a wrong conclusion. In fact, for Aβ40 and by extension other IDPs, RMSD values happen to be useless for judging whether an MD simulation has reached convergence.

#### Number of clusters

Next, we calculated the number of conformational Aβ40 clusters using the clustering method of Daura et al. and an RMSD cutoff of 1.0 nm as a function of simulation time. The results in Figure 1 show that for almost all FFs more than 10 µs of MD sampling is needed before the curves for this quantity converge. But for several of the FFs even beyond 20 µs new conformations are still sampled. Only with a03ws, a99SB*-ILDN/TIP3P and C22*/TIP3P no new conformations were found in the last ∼20 µs of the MD trajectories. Another difference between these and the other FFs is that the total number of clusters is considerably smaller. With a99SB*-ILDN/TIP3P, less than 5 conformational clusters were identified, whereas with C36m more than 30 clusters were sampled. Thus, the different FFs predict different degrees of Aβ40 structural flexibility and the FFs associated with higher conformational diversity were generally observed to require simulation times longer than 20 µs for attaining convergence. It should be noted that due to the use of relatively large RMSD cutoff (1.0 nm), the different clusters considerably vary from each other structurally. Put differently, this implies that transitions from one cluster to another involve non-negligible conformational changes.

#### Radius of gyration

The assessment of *R*_gyr_ (Figure S1 in the Supporting Information) reveals that this quantity equilibrates quickly and in most cases did not considerably change after 10 µs. This applies especially to a99SB-ILDN/TIP4P-D and a99SB-UCB for which more than 20 µs of simulation time is needed for the number of sampled clusters to converge; the two corresponding *R*_gyr_ distributions however do not change after a simulation time of 10 µs. It is only with a03ws that a considerable change in the *R*_gyr_ distribution occur after 10 µs. It changes from a broad distribution with a maximum value of ∼1.5 nm to a narrow distribution with a distinct peak at around ∼1.1 nm and a flat distribution with rather low probabilities for larger *R*_gyr_ values. Thus, a considerable conformational transition must have happened shortly after 10 µs, leading to a conformation considerably different and more compact to all previously sampled conformations. Figure 1 shows that once this structure was identified, no further new structures were sampled as the number of clusters did not rise after ∼11 µs in the case of a03ws. Comparison of the *R*_gyr_ distributions obtained for the different FFs reveals that quite different structural ensembles are produced: only compact structures with a99SB*-ILDN/TIP3P, compact and also elongated structures with a03ws, a99SB-disp, C22*/TIP3P and C36m, and mostly elongated structures with a99SB-ILDN/TIP4P-D and a99SB-UCB.

#### Secondary structure

Further information on the structural preferences obtained for the different FFs is available by analyzing the secondary structure. Figure 2 shows the evolution of the amounts of α-helix and β-sheet found in Aβ40, while in Figure S2 the time averages for the secondary structure elements can be seen. After 10 µs these time averages are largely converged apart for a03ws. In the latter case, a gradual increase in helix is observed till the end of the simulation, whereas for the other FFs a slight increase in β-sheet is seen for sampling times above 10 µs. The propensity for β-sheet formation differs with the FFs: a99SB-ILDN/TIP4P-D and a99SB-UCB predict a low amount of β-sheet close to zero, a medium amount of β-sheet is found for a03ws, a99SB-disp, C22*/TIP3P and C36m with values of ∼10–15%, while a99SB*-ILDN/TIP3P generates a conformational ensemble with more than 20% of the Aβ40 residues adopting a β-sheet structure. The tendency of Aβ40 to adopt helical structures is close to zero for a99SB-ILDN/TIP4P-D, a99SB-UCB, and C36m, whereas with a03ws almost 20% helical content is observed at the end of the simulations. These differences in secondary structure preferences correlate well with the different *R*_gyr_ distributions. For example, a high amount of helix and/or β-sheet lead to compact structures as observed for a03ws and a99SB*-ILDN/TIP3P, whereas low amounts of helix and β-sheet imply elongated structures as is the case for a99SB-ILDN/TIP4P-D and a99SB-UCB.

It is interesting to not only assess time averages for the secondary structure but also its evolution. Figure 2 reveals that within 10 µs – the time window chosen for averaging – considerable changes in secondary structure can occur. This applies to all FFs yet to different extents. The smallest changes occur for a99SB-ILDN/TIP4P-D and a99SB-UCB, i.e., the two FFs which generally predict a low tendency for Aβ40 to adopt a helical or a β-sheet conformation. Another extreme is C36m for which huge changes in the amount of β-sheet are observed after 10 µs. It is also interesting to correlate the secondary structure changes to the number of clusters, revealing that even within the same cluster considerable secondary structure changes can occur. This is best seen for the C36m simulation at ∼5–20 µs. During this period, the number of clusters is constant, while the amount of β-sheet formed within Aβ40 first varies widely between 0 and 20%, then increases to more than 40% between 13 and 14 µs, followed by a decrease until no more β-sheet is present at around 18 µs.

### 3.2 Convergence checks based on Markov state models

#### Sample density in TIC space

An ultimate test to determine whether or not the MD data has converged is possible by calculating an MSM; this usually requires the performance of a Chapman-Kolmogorov test for the level of agreement between the MSM predicted dynamics and the actual protein dynamics. Several steps in the construction of an MSM involve dimension reduction and often implies TICA as used in this study. The projection of the MD data along the first two TICs can be seen in Figure 3. To assess the evolution of the Aβ40 conformations in TIC space, we projected the data from 10 µs time windows. For each of the FFs one can see that different conformational spaces are sampled for the different time windows. Nonetheless, for all FFs but a99SB*-ILDN/TIP3P the explored spaces partially overlap. To verify that no no new structures are sampled for longer simulation times, we extended the simulations for another 5 µs for a99SB-disp and C36m. The projection in TIC space (Figure S3) shows that indeed no new conformations are acquired.

**Figure 3:**
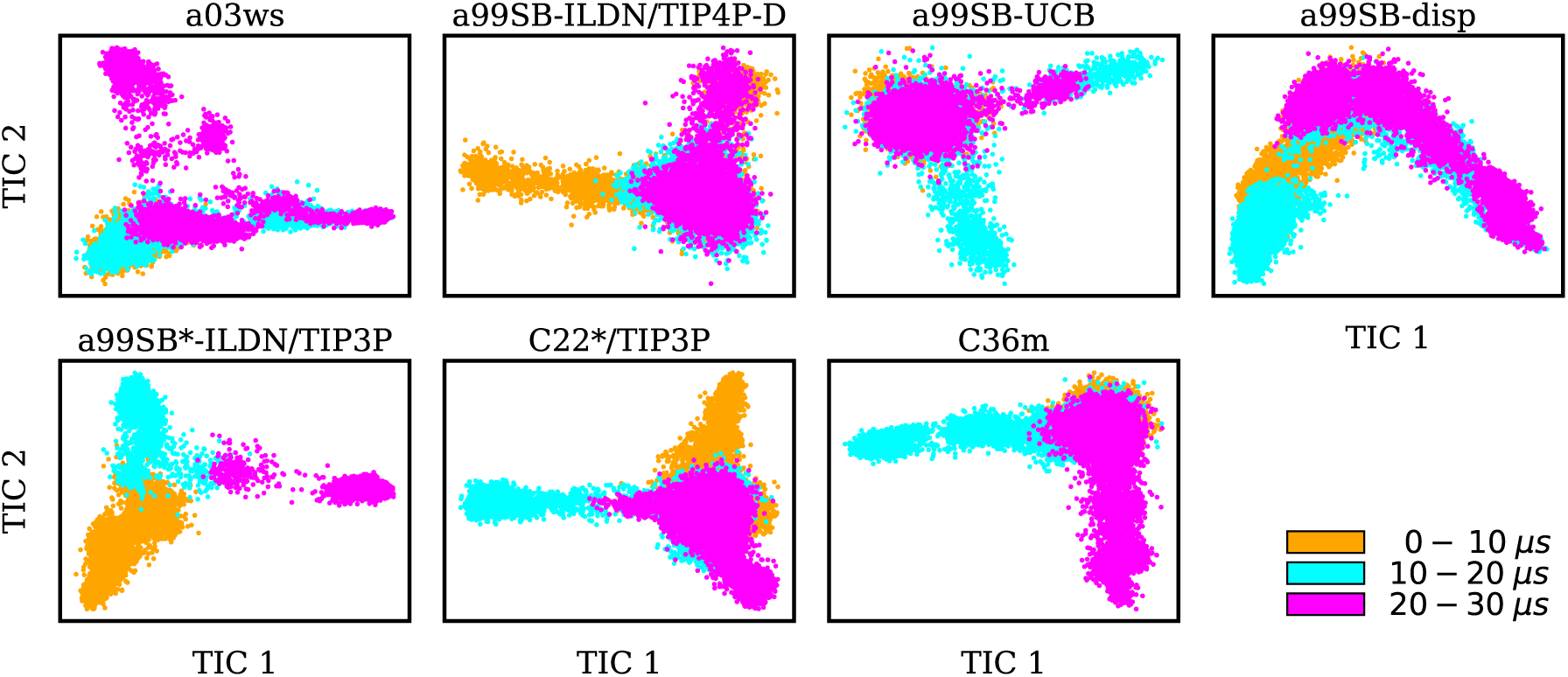
Sample densities for different time windows of the trajectories (0–10 µs: yellow, 10–20 µs: cyan, 20–30 µs: magenta) projected along the first two TICA coordinates (TICs) for the different force fields (labels on the top of the panels). It should be noted that the TICs are different between the force fields.

Taking C36m as an example for a detailed analysis of the sample density, one finds that within the first 10 µs Aβ40 largely remained within the same region of the TIC space. Between 10 and 20 µs it explored new conformations (along TIC 1), which, as revealed by the analysis of the secondary structure, resulted from first a build-up of a β-sheet conformation, followed by its destruction. It should be emphasized that TICA identifies the slowest modes in the feature space and not, unlike principal component analysis, the modes of largest motions. TICA thus confirms the result from Figure 2 that the formation and also disassembly of β-sheets is a slow process in Aβ40, requiring several µs of MD sampling (not only with C36m, but also with.e.g., a99SB-UCB and a99SB-disp). Figure 3 further shows that after 20 µs of MD with C36m a novel conformational transition, evolving along TIC 2, is explored. Comparison with Figure 2 reveals that this process again involves β-sheet formation, followed by its disassembly. In the additional 5 µs trajectory, Aβ40 remained in the energy well corresponding to this β-sheet structure (Figure S3).

#### Markov state models

The final coarse-grained MSMs (also called HMMs) are shown in Figure 4. In the case of a99SB*-ILDN/TIP3P the convergence of the MD data was not sufficient to allow for the construction of an MSM. Thus, only for the other six FFs Markov state models are shown. The MSMs for a99SB-ILDN/TIP4P-D and a99SB-UCB are similar to each other as are their previously discussed observables. These two MSMs are dominated by a single macrostate with a population of 95% (state 3 in both MSMs) and two further low-populated states. To characterize the macrostates we calculated the inter-residue contacts for all MD frames assigned to each of the states and averaged the contacts per mascrostate (Figures S4–S9). The contact maps of states 3 for these two FFs (Figures S5 and S6) involve almost no contacts between residues which are not in neighborhood of each other in the primary structure, i.e., these conformations are mainly elongated structures with large *R*_gyr_ values and with low amounts of helix and β-sheet.

**Figure 4:**
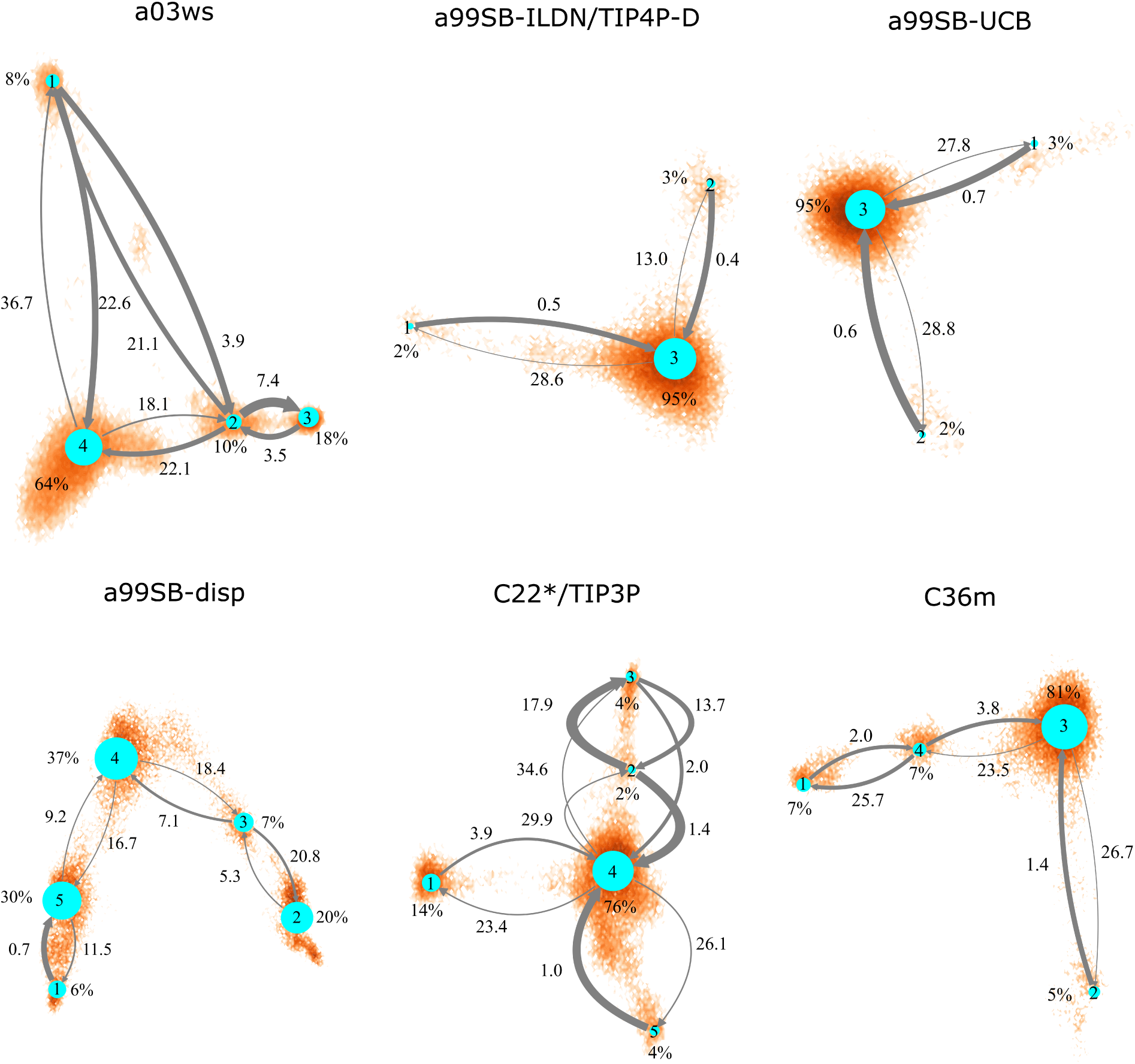
Coarse-grained Markov state models for the different force fields (labels above the networks). The size of the network nodes reflects the population of the underlying state (given in % next to the nodes), whereas the thickness of the arrows corresponds to the transition probability. The mean first passage times (in µs) are written next to the arrows. The orientation of the MSMs corresponds to the projection of the MD data along the first two TICs in Figure 3.

The MSM for C36m also involves a dominant state corresponding to a stretched structure with no noteworthy contacts between distant residues (state 3, 81% population). However, with C36m also more compact structures with β-sheets are adopted, yielding MSM states 1, 2 and 4 (19% population in total). These three states exhibit a similar antiparallel β-sheet (Figure S9). In states 1 and 4 this involves a β-hairpin centered at residues G25/S26. The β-sheet in state 4 is on-pathway to that of state 1 (coming from state 3) but is shorter than the fully-formed β-sheet of state 4. The latter involves up to 8 or 9 residues per strand and thus reaches up to residue 15 towards the N-terminus and residue 35 towards the C-termins. This corresponds to the maximum count of 16 to 17 residues that adopt a β-sheet conformation between 13 and 14 µs of the MD simulation (Figure 2). Another characteristic of the β-sheet in states 1 and 4 is the salt bridge present between D23 an K28, giving rise to the strongest inter-residue contact in either state.

Similar β-sheet formation is observed with C22*/TIP3P and a99SB-disp, yet the β-sheets are less pronounced (a99SB-disp, Figure S7) and may also occur in the N-terminal half of the peptide (C22*/TIP3P, Figure S8). Furthermore, the MSMs for these two FFs do not feature a state representing an elongated Aβ40 structure. Instead, more structure formation is seen which also involves helical elements seen in the C-terminal half of the peptide in states 2 and 3 of the MSM obtained with C22*/TIP3P. These two states, however, are only characterized by a low population (2 and 4%, respectively), while the most populated state observed with this FF is associated with a low interpeptide contacts probability (state 4, 76% population) close to an elongated structure. The four states of the MSM for a03ws all feature a β-sheet in the first 10 N-terminal residues, but differ from each other in their contacts in the rest of the peptide (Figure S4). Helices of various lengths and locations along the sequence are presents in states 1–3, while the most populated state 4 (64% population) is without noteworthy inter-residue contacts beyond residue 10, thus representing a mainly elongated structure.

Comparison between Figures 3 and 4 (the MSMs from Figure 4 can be superimposed onto the sample density in Figure 3 as the same TIC space is used for their projection) show that several of the metastable MSM states were only sampled in the last 10 µs. This holds for instance true for state 1 of the MSM with a03ws and also state 2 of the MSM with C36m. The calculation of the mean first passage times (MFPTs) for the transitions between the metastable states confirms that the slowest transitions require more than 20 µs for them to occur. Such slow transitions are found in all of the MSMs shown in Figure 4. As this work aims to provide an answer to the question how much MD sampling is needed before the equilibrium distribution of Aβis reached, based on the MFPTs the answer would be that 30 µs are required if a single MD trajectory is being collected. However, it should be mentioned MSMs can integrate the data of short MD simulations and thus effectively parallelize the computational effort. However, the rare transitions still need to be spontaneously sampled in the data.

### 3.3 Validation of simulated Aβ40 ensembles based on spectroscopic data

In order to validate the simulation results before further discussing them, we compare the structural ensembles obtained for Aβ40 with the corresponding information deduced from spectroscopic data, which is NMR chemical shifts and *J* -couplings^59^ as well as FRET efficiencies.^35^ In addition, we also compare the simulated and experimentally determined dynamics of Aβ40.

#### Chemical shifts

We calculated the NMR chemical shifts of the carbonyl C′and C_*α*_ atoms for all MD generated conformations and provide the averages over the 30-µs MD simulations in Figure S10. The agreement between the measured and calculated C_*α*_ chemical shifts is generally good for all FFs as judged by the RMSD between these data sets (Table 1). The smallest RMSD is found for a99SB-ILDN/TIP4P-D with a value of 0.59 ppm, while the largest values of 0.74 and 0.76 ppm result from a03ws and a99SB*-ILDN/TIP3P, respectively. A similar picture emerges if one compares the C′-chemical shifts. By far the largest deviation is found for a03ws with a value of 1.03 ppm, whereas a99SB-UCB and a99SB-disp lead to the smallest RMSDs of 0.93 ppm. A preliminary conclusion is that a03ws is the least agreeable with the employed NMR chemical shifts data. The comparison of the chemical shifts for each residue shows that the larger deviation in comparison to the other FFs results from chemical shifts on the N-terminal side, especially for Y10 and D11. The calculated C_*α*_- and C′-chemical shifts are higher than the experimental values, indicating an overestimation for α-helix formation in the simulation with a03ws.^60^

**Table 1:** Simulated properties of Aβ40 sampled with different force fields

| Quantity | Force fields |  |  |  |  |  |  |
| --- | --- | --- | --- | --- | --- | --- | --- |
|  | a03ws | a99SB-ILDN/TIP4P-D | a99SB-UCB | a99SB-disp | a99SB*-<br>ILDN/TIP3P | C22*/TIP3P | C36m |
| RMSD<sub>exp</sub> C$_{\alpha}$<br>shift (ppm) | 0.738 | 0.589 | 0.641 | 0.631 | 0.759 | 0.688 | 0.634 |
| RMSD<sub>RC</sub> C$_{\alpha}$<br>shift (ppm)<sup>#</sup> | 0.838 | 0.658 | 0.700 | 0.690 | 0.791 | 0.756 | 0.686 |
| RMSD<sub>exp</sub> C'<br>shift (ppm) | 1.025 | 0.964 | 0.927 | 0.929 | 0.984 | 0.961 | 0.978 |
| RMSD<sub>RC</sub> C'<br>shift (ppm)<sup>#</sup> | 0.766 | 0.672 | 0.658 | 0.653 | 0.734 | 0.707 | 0.745 |
| $\chi^2 J$-coupling | 7.03 | 3.28 | 1.94 | 4.50 | 6.33 | 2.50 | 3.05 |
| $\langle R_{ee} \rangle$ (nm) | $3.00 \pm 1.80$ | $4.52 \pm 1.70$ | $4.81 \pm 1.90$ | $3.61 \pm 1.40$ | $2.53 \pm 0.90$ | $3.04 \pm 1.20$ | $3.52 \pm 1.7$ |
| $\langle E_{\text{FRET}} \rangle$ | $0.83 \pm 0.26$ | $0.65 \pm 0.31$ | $0.59 \pm 0.33$ | $0.81 \pm 0.23$ | $0.96 \pm 0.06$ | $0.88 \pm 0.17$ | $0.80 \pm 0.27$ |
| $\langle R_{\text{gyr}} \rangle$ (nm) | $1.31 \pm 0.30$ | $1.65 \pm 0.30$ | $1.76 \pm 0.40$ | $1.38 \pm 0.30$ | $1.11 \pm 0.08$ | $1.24 \pm 0.20$ | $1.45 \pm 0.3$ |

<sup>#</sup> For the calculation of the RMSD with respect to the RC chemical shifts, the RC values from ref 61 were used.

Bax and co-workers concluded from their NMR studies that both Aβ40 and Aβ42 predominantly sample random coil (RC) structures. To better estimate how much the simulated ensembles deviate from random coil, we also calculated the RMSDs between the calculated and RC chemical shifts (Table 1). For the latter we used the data set by Wang and Jardetzky,^61^ which is somewhat different from the RC shifts applied in ref 59. The RMSD rankings with respect to the experimental chemical shifts and with respect to the RC values are very similar. In both cases the largest deviations are found for a03ws, followed by a99SB*-ILDN/TIP3P. Based on Figure 2 it can be concluded that the latter FF overestimates the tendency to form β-sheets. Closest to RC structures are the structural ensembles generated with a99SB-ILDN/TIP4P-D, a99SB-UCB and a99SB-disp, followed by C22*/TIP3P. These findings agree with our conclusions drawn from the MSMs in conjunction with an inter-residue contact analysis for the macrostates. Interestingly, C36m, which was purposefully developed for IDPs,^31^ leads to less RC in the structures of Aβ40, which was also observed in the contact maps of the MSM states as three of the four MSM states feature rather high β-sheet formation (Figure S9). We further find that for the C′-atoms the RMSDs from the RC chemical shifts are smaller than the RMSDs from the experimental chemical shifts. On the other hand, the RMSD between the experimental C′-chemical shifts and the corresponding RC values is even smaller with a value of 0.56 ppm. This observed divergence in how both the predicted and experimental chemical shifts differ from the RC values necessitates an in-depth analysis.

For all FFs the largest deviation between simulation and experiment is found for V24, K28, I32 and V36 (Figure S11A). For these residues, the measured chemical shifts are considerably larger than the calculated ones. Comparison of the RC and measured C′-chemical shifts reveals similar deviations for these four residues (Figure 1 in ref 59), while the calculated values are relatively close to the RC values (Figure S11B). The experimental values suggest that these residues have a tendency to adopt a helical conformation. Under consideration of all of the up to 12 measured NMR parameters this tendency was confirmed for V24 and K28 by the MERA program (Maximum Entropy Ramachandran map Analysis from NMR data)^62,63^ that was developed by the Bax lab and applied to the Aβ40/42 NMR data.^59^ For I32 the deviation from the RC chemical shift is small if one takes the RC values and correction factors of Poulsen and co-workers,^64,65^ while for V36 a preference for RC is found if all other NMR data measured for that residue are considered. For the region A30–I32 all the predicted C′-chemical shifts are smaller than the experimental values for all FFs. The smallest deviation is found with a99SB-UCB, which also explains the overall smaller RMSD from experiment found for this FF. Comparison with the RC values shows that with a99SB-UCB this region is in a RC state, while the other FFs sample to some degree a β-conformation here.^60^

Another region that needs attention is V17–A21 forming the central hydrophobic core (CHC), which by many studies is proposed to adopt a β-conformation and play a central role during amyloid aggregation of Aβ.^8^ There is almost no deviation between simulation and experiment for V18 and F19, while for the other residues and also the neighboring K16 and E22 the simulations lead to smaller C′-chemical shifts than in experiment. The predicted values are also smaller than the RC chemical shifts, indicating that this region tends to adopt a β-conformation in the simulations, which was also seen in several of the MSM states for most of the FFs. The NMR data, on the other hand, indicate β-strand formation only for V18–F20, which follows from the comparison between the RC and experimental chemical shifts and also from the MERA anaylsis that considers the other NMR data.^59^ However, Bax and co-workers excluded β-sheet formation for that region as from their NMR data no matching set of residues with which to pair these residues in a β-sheet could be identified. In the simulations these residues usually pair with I30–L34 (Figures S4–S9), which, as already discussed above, also tend to be in a β-conformation.

#### *J* -couplings

Bax and co-workers did not only record chemical shifts for Aβ40 and Aβ42, but also three-bond *J* -couplings, including ^3^*J*_HNH*α*_ couplings^59^ that are related to the backbone torsion angles ϕ by the empirically parametrized Karplus equation. We use these experimental values to further validate the Aβ40 ensembles generated by MD simulations with different FFs. The comparison between simulation and experiment for the C^*I*^ and C_*α*_ chemical shifts led to the conclusion that a99SB-ILDN/TIP4P-D, a99SB-UCB and a99SB-disp produce Aβ40 structural ensembles best in agreement with the NMR chemical shift data, while a03ws and a99SB*-ILDN/TIP3P fail to do so. A closer inspection of the C^*I*^ chemical shifts revealed that a99SB-UCB is in particular able to reproduce Aβ40’s tendency to adopt a RC state. Comparison for the ^3^*J*_HNH*α*_ couplings confirms that this FF is superior to the other FFs in modeling Aβ40 as a *χ*^2^ value of only 1.94 is obtained (Table 1). The *χ*^2^ values for a03ws and a99SB*-ILDN/TIP3P of 7.03 and 6.33, respectively, also confirm the previous conclusion, which is that these two FFs do not yield good structural ensembles for Aβ40.

For the remaining four FFs the assessment is somewhat more complicated. The usage of a99SB-ILDN/TIP4P-D can be recommended as *χ*^2^ for the ^3^*J*_HNH*α*_ couplings is also still considerably small with a value of 3.28; though a99SB-UCB performs clearly better. Interestingly, a99SB-disp, which was developed for IDPs (but also folded proteins) and claimed to be one of the best FFs for Aβ40,^32^ does not yield ^3^*J*_HNH*α*_ couplings in agreement with experiment. The *χ*^2^ value is 4.50 and comparison of the *J* -couplings on a per residue basis (Figure 5) shows that for most residues these values are smaller than the experimental *J* -couplings, only reaching values of ∼ 6 or less. An exemption to this is the region K16–E22 where the CHC residues V18–F20 in particular demonstrate a tendency to adopt β-conformation, which is also correctly reflected by the *J* -couplings. Here, a very good agreement with the experimental values is obtained for a99SB-disp. For the other residues, on the other hand, the average ϕ values must be too small. To validate this conclusion we analyzed the Ramachandran angles of I31 and I32 as sampled in the MD simulations in detail (Figure S12). From experiment, ^3^*J*_HNH*α*_ couplings above 7 Hz were obtained, indicating a considerable population of extended structures, which is supported by MERA.^59^ Figure S12 shows that with a99SB-disp only few structures with ϕ*<* −90^°^ were sampled for I31 and I32.

**Figure 5:**
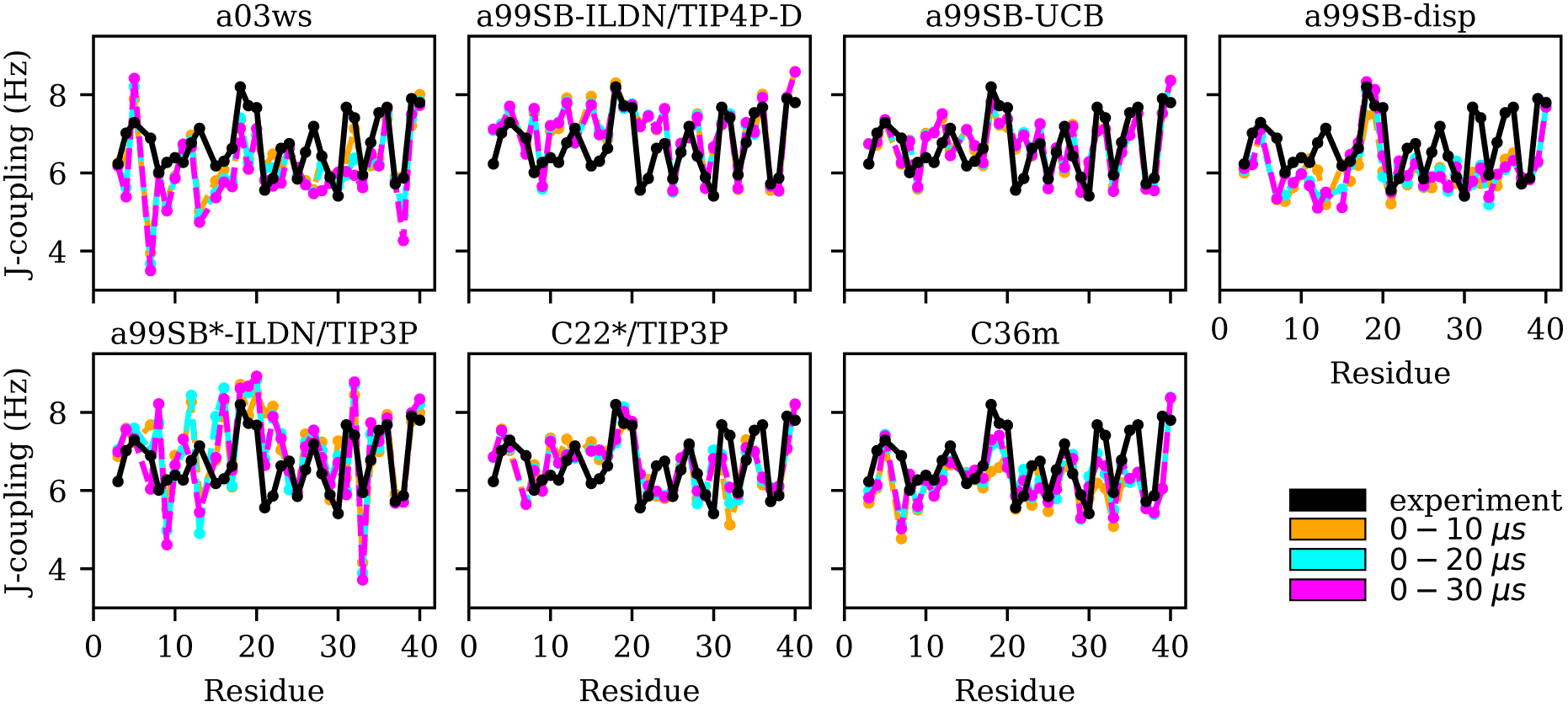
Experimental (black) and simulated (0–10 µs: yellow, 0–20 µs: cyan, 0–30 µs: magenta) ^3^*J*_HNH*α*_ couplings for each Aβ40 residue for the different force fields (labels on the top of the panels).

Instead, polyproline II (PPII) conformations were preferentially adopted, which explain the low *J* -couplings. With a99SB-ILDN/TIP4P-D and a99SB-UCB, on the other hand, which both led to a good agreement for the *J* -couplings of I31 and I32, a considerable amount of conformations with ϕ *<* −120^°^ were sampled, which includes extended conformations from the β-basin as well from the type I β-turn region. Interestingly, with these two force fields the largest variability of conformations was adopted. Basically all allowed regions of the Ramachandran space, including the α_L_ region were sampled, which agrees with the notion that random coil is not a particular structure but a fast fluctuation between all possible ϕ/ ψ combinations.

For the two Charmm FFs under consideration, the situation is contrary to that of a99SB-disp. For C22*/TIP3P and C36m the agreement with NMR chemical shifts is largely insufficient, while the *χ*^2^ values for the ^3^*J*_HNH*α*_ couplings are satisfactory. Interestingly, the FF not improved for IDPs, i.e., C22*/TIP3P performs somewhat better for both chemical shifts and *J* -couplings compared to C36m. However, the disagreement between both FFs is limited to a few residues, such as D7, M35 and V36 where C36m performs worse for the *J* -couplings, while for most of the remaining residues similar NMR observables are predicted. For M35 and V36, C36m sampled a high amount of PPII and α_R_ structures, for M35 also α_L_ (based on the corresponding Ramachandran plots, not shown), leading to *J* -couplings below those found from experiment, which however, as Figure 5 shows, increased for sampling times above 10 µs. In general, we observed that it was only beyond 10 µs simulation times that MD convergence for the *J* -couplings was obtained. This conclusion excludes a99SB-ILDN/TIP4P-D and a99SB-UCB, for which converged *J* -couplings were already obtained within 10 µs.

#### End-to-end distance and hydrodynamic radius

Another experimental observable that is available for Aβ40 was derived from 2D FRET data that have been reported by Meng et al.^35^ They determined the end-to-end distance, *R*_ee_, of Aβ40 from the analysis of the FRET efficiency between the donor Alexa 488 and the acceptor Alexa 647, which were attached at the termini. To this end, an unnatural amino acid, 4-acetylphenylalanine and a cysteine residue were first introduced at the N- and C-terminus of Aβ40, respectively. An average FRET efficiency of ∼0.6 was obtained. According to eq (2) this corresponds to a distance of 4.85 nm between donor and acceptor, called 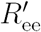 here. To account for the size of the fluorophores, eq (3) is applied, yielding an average end-to-end distance of∼4.3 nm for Aβ40. The values for *R*_ee_ and the FRET efficiencies in Table 1 show that a99SB-ILDN/TIP4P-D and a99SB-UCB perform very well in reproducing these observables. While C36m and a99SB-disp are next in performance, they however clearly underestimate *R*_ee_, leading to overestimated FRET efficiencies of 0.80 and 0.81, respectively. Even more compact Aβ40 structures are produced by C22*/TIP3P and a99SB*-ILDN/TIP3, with the latter FF leading to average FRET efficiencies of nearly 1 (*E*_FRET_ = 0.96). For a03ws the situation is somewhat more complicated. The distributions of *R*_ee_ averaged over different times (Figure 6) shows that for most FFs this quantity converged within 10 µs. Exceptions are a03ws and a99SB*-ILDN/TIP3P. With the former FF, the end-to-end distance became smaller with time, with a pronounced peak that developed at Ree ∼1.2 nm. With a99SB*-ILDN/TIP3P, on the other hand, the peak at *R*_ee_ ∼0.9 nm became less important with time, while more extended structures were adopted. The evolution of *R*_ee_ over the whole trajectory (Figure S13) reveals that with a03ws large fluctuations with respect to the end-to-end distance occurred, involving the formation of a compact conformation with especially low *R*_ee_ values in which Aβ40 was trapped between 16 and 25 µs. Representative conformations from this trapped state are shown in Figure S14. They are characterized by the presence of helices along the sequence from residue Y10 onward, corresponding to MSM states 1, 2 or 3. This finding is surprising as the increase in van der Waals interactions between protein and water as done in the development of a03ws was thought to avoid such overly compact protein states,^30^ yet the current 30 µs MD simulation shows that the helical propensity present in the FF predecessor a03^66^ overrules this modification if simulated long enough. With the other FFs such trapping is not seen; instead, fast fluctuations in *R*_ee_ are sampled, with C36m showing the largest and C22*/TIP3P and a99SB*-ILDN/TIP3 the smallest fluctuations.

**Figure 6:**
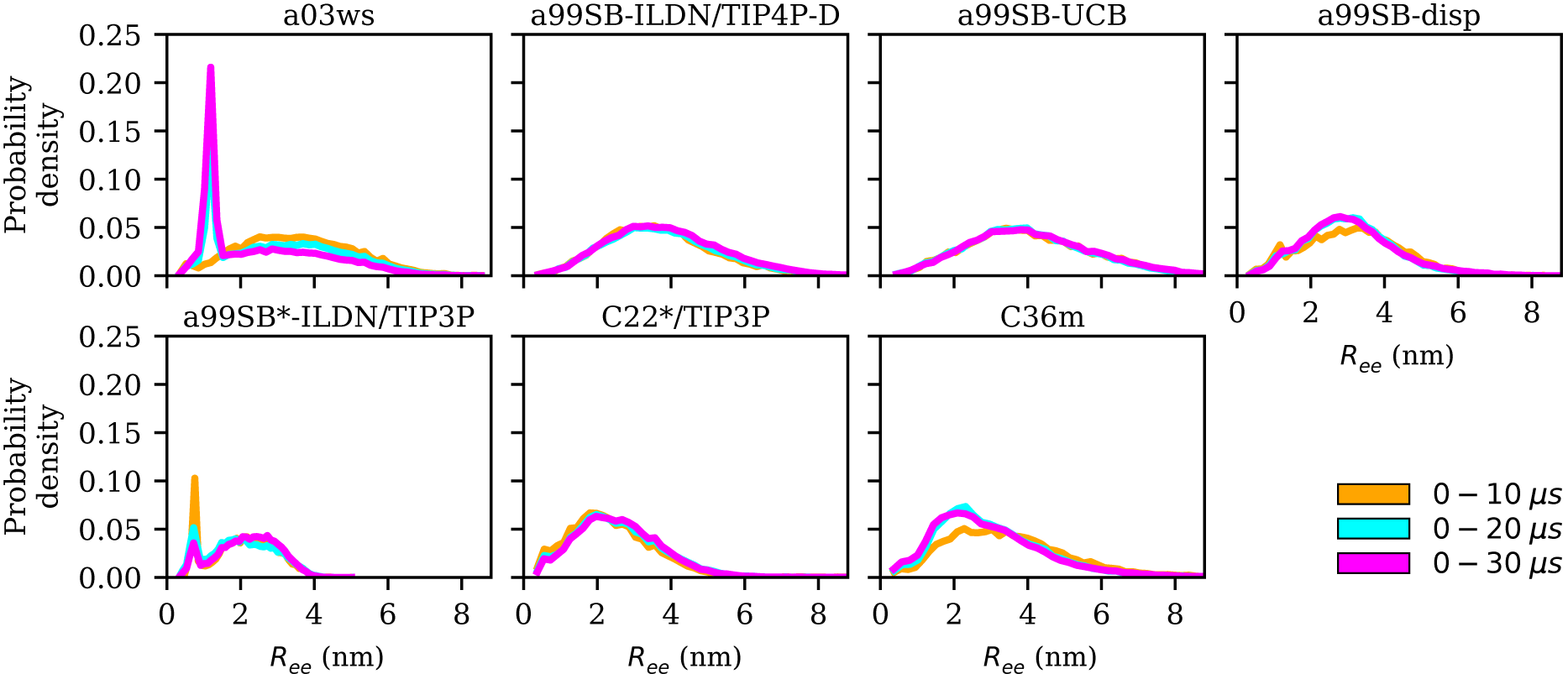
Distribution of the end-to-end distance *R*_ee_ for increasing trajectory lengths (0– 10 µs: yellow, 0–20 µs: cyan, 0–30 µs: magenta) for the different force fields (labels on the top of the panels).

While the agreement between average FRET efficiencies determined experimentally and those derived from the simulations with a99SB-ILDN/TIP4P-D and a99SB-UCB is very good, the same cannot be said for the distribution of the FRET efficiencies. In experiments this distribution assumes a Gaussian shape around the average value,^35^ while we find highly skewed distributions with the maximum for FRET efficiencies being close to one (Figure S15). The same observation was made in two previous simulation studies where, as observed here with a99SB-ILDN/TIP4P-D and a99SB-UCB, the agreement with the average *E*_FRET_ value was associated with a considerable amount of structures with *R*_ee_ values above 6 nm.^35,56^ Further simulations are needed to identify the source of this disagreement. There, effects of the FRET labels including the extra amino acids added at the termini of Aβ should be explicitly considered as well as the orientation of donor and acceptor with respect to each other be accounted for when determining *E*_FRET_.

Another quantity closely related to *R*_ee_ is the hydrodynamic radius, *R*_hyd_, which was determined as 1.6 nm for Aβ40 at 298 K using size exclusion chromatography (SEC) and NMR diffusion experiments.^67^ In general, the hydrodynamic radius is closely related to the radius of gyration. For IDPs, a releationship between these two quantities was derived that explicitly takes the chain-length dependency into account.^68^ Using eq (7) of ref 68 one obtains for an IDP with 40 amino acids that a hydrodynamic radius of 1.6 nm corresponds to a radius of gyration of 1.2–1.3 nm. This *R*_gyr_ value is best reproduced by a03ws and C22*/TIP3P, followed by a99SB-disp and C36m. With a99SB*-ILDN/TIP3P *R*_gyr_ is underestimated, while a99SB-ILDN/TIP4P-D and a99SB-UCB lead to *R*_gyr_ values clearly above the experimental prediction. This observation is mostly in contrast to the conclusions drawn from the calculation of *E*_FRET_ since for *R*_ee_ the best agreement was identified for a99SB-ILDN/TIP4P-D and a99SB-UCB. Only for a99SB*-ILDN/TIP3P both *R*_ee_ and *R*_gyr_ agree with each other, both being too small compared to experiment as a result of too compact Aβ40 structures being sampled with this FF. Since for all other quantities a99SB-ILDN/TIP4P-D and a99SB-UCB produced the best agreement with experiment, we decided to disregard the assessment based on *R*_gyr_, especially since it contradicts the *R*_ee_ results.

### 3.4 Validation of simulated Aβ40 kinetics based on spectroscopic data

The kinetic analysis of the MD data using MSMs allows to further assess the accuracy of the simulation data based on time scales that were reported for Aβ motions. From the FRET study mentioned above^35^ it was concluded that Aβ40 and Aβ42 exhibit no conformational dynamics exceeding 1 µs and that the end-to-end distance relaxation time is ∼35 ns, which was determined by nanosecond fluorescence correlation spectroscopy (nsFCS). The upper limit of 1 µs for internal motions agrees with the findings from fluorescence measurements using the method of Trp-Cys contact quenching, which revealed a time scale of ∼1 µs for intramolecular reorientation or diffusion for Aβ40.^36^ With NMR spin relaxation measurements the faster motions of Aβ40 were studied, from which a timescale of ∼5 ns was determined for segmental motions, which can reach up to ∼10 ns for selected residues.^69^ The focus of MSMs is the identification of slow, memoryless motions. Thus, for the current comparison of the Aβ40 kinetics as determined by experiment and MD simulations, the upper limit of 1 µs for the slowest motion is of interest to us. All FFs with conformational transitions exceeding this time scale can be rendered as inadequate for modeling the kinetics of Aβ. Here it should be emphasized that the MFPTs discussed above refer to well-defined time scales between specified states, which are not the same as the relaxation times probed by the different spectroscopic techniques. For this, the implied time scales (ITSs) underlying the constructed MSMs are better suited. The implied time scales reflect how quickly any initial state vector converges towards the equilibrium state vector in an MSM and are thus comparable to relaxation times monitored experimentally. The MFPTs, on the other hand, indicate the time it takes to transition from one equilibrium state vector to another one.

This can become considerably larger than the ITSs, especially for transitions into equilibrium states with very small probabilities, which can be seen from the MFPTs in Figure 4. Thus, we concentrate on the ITSs here.

The plots of the ITSs against the lag times of the MSMs (Figure S16) show that in the case of a03ws, a99SB-disp, C22*/TIP3P and C36m they clearly exceed 1 µs. Some of these FFs even reach time scales for the slowest motion of more than 10 µs (a03ws and a99SB-disp). It should be noted that for a99SB*-ILDN/TIP3P no results for the implied time scales can be shown as with this FF the slowest dynamics of Aβ40 reached the length of the MD trajectory, i.e., 30 µs. Thus, a99SB-ILDN/TIP4P-D and a99SB-UCB are the only two FFs which agree with the experimental finding that the slowest intramolecular Aβ40 dynamics takes places within 1 µs. The exact time scales are 500 ns for a99SB-ILDN/TIP4P-D and 700 ns for a99SB-UCB.

## 4 Discussion

### 4.1 Which force fields are suitable for modeling Aβ?

Based on the detailed comparison with NMR and FRET data we can conclude that a99SB-UCB produced an Aβ40 conformational ensemble that is best in agreement with experiment. The second best FF is a99SB-ILDN/TIP4P-D. Like a99SB-UCB it produces extended Aβ40 conformations in agreement with FRET. Also the NMR chemical shift data are of similar quality, yet the ^3^*J*_HNH*α*_ couplings are less in agreement. The performance of the two Charmm FFs, C22*/TIP3P and C36m, is similar to each other. While the older of these two FFs performs better in reproducing the NMR data, the IDP corrected FF yields, on average, less compact Aβ40 conformations. However, also with C36m overly compact structures with too much β-sheet content are still sampled. The overall still acceptable agreement with experimental findings is realized by the extended structures that are temporarily adopted with this FF. It would be interesting to test C36m with a water model with more favourable LJ interactions between protein and water. This approach improved the performance of C36m for some of the IDPs that were studied by the developers of C36m.^31^ It should be noted that in our previous FF benchmark for Aβ42 we had identified C22*/TIP4P-Ew as the best FF for Aβ42.^17^ However, this benchmark did not include any of the FFs recently developed for IDPs since it was performed prior to their development. Nonetheless, even though C36m was explicitly developed for IDPs (by refining backbone parameters),^31^ it does not lead to a better performance than the standard C22*/TIP3P force field.

Another FF that was developed for IDPs (but also for folded proteins) is a99SB-disp.^32^ Its developers claimed that it is one of the best FFs for Aβ40. However, our analysis revealed that it fails to produce Aβ40 structures in agreement with ^3^*J*_HNH*α*_ coupling data determined by NMR spectroscopy. The underlying reasons for this is that a99SB-disp shows a preference for PPII conformations, which consequentially leads to an underestimation of the ^3^*J*_HNH*α*_ couplings for many of the Aβ40 residues. Therefore, the use of this FF for modeling Aβ40 is not recommended. Least suitable for modeling Aβ are a03ws and a99SB*-ILDN/TIP3P, which should not be applied to Aβ. After 16 µs of MD with a03ws, Aβ40 became trapped in a highly compact, helical state (Figure S14), which is unsupported by any available experimental data. With a99SB*-ILDN/TIP3P as well, overly compact conformations are found, which in this case result from excessive β-sheet structures.

One question however remains: can one understand the basis for the superior performance observed with a99SB-ILDN/TIP4P-D and a99SB-UCB compared to the other FFs? It is proper to directly recognize our inability to comment on how different modifications of the force fields not covered in the current work would influence their performance. For instance, we cannot say for sure whether adjusting the ψdihedral parameters in a99SB* (that is, compared to a99SB) played a role in the performance performance of a99SB*-ILDN/TIP3P since we neither have a 30 *µ* simulation with a99SB-ILDN/TIP3P nor with a99SB*-ILDN/TIP4P-D as references. However, based on the results from our previous benchmark, ^17^ where a99SB/TIP4P-Ew and a99SB*-ILDN/TIP4P-Ew produced almost identical results for Aβ42 (i.e., the adjustment of dihedral parameters made no difference), the conclusion is that the main reason for a99SB*-ILDN/TIP3P being an inadequate FF choice for Aβ is the water model. And the same holds true for the good performance of a99SB-ILDN/TIP4P-D and a99SB-UCB. The latter FF is based on a99SB/TIP4P-Ew with a modified C-N-C*α*-Cβ dihedral angle (ϕ′) potential, which led to improved conformational ensembles for a number of model peptides,^33^ and optimized LJ interactions between protein and water, that yielded better agreement with experimental solvation free energies for 47 small molecules that incorporated all of the chemical functionalities of standard protein side chains and backbone groups.^34^ The conclusion that the revised protein–water interactions are a key ingredient is further supported by the fact that a99SB/TIP4P-Ew produced too compact and too much structured Aβ42 conformations in our previous benchmark.^17^

The same arguments apply to a99SB-ILDN/TIP4P-D. The water model TIP4P-D is based on TIP4P/2005, but compared to that features a significantly higher dispersion coefficient C6, which, like in a99SB-UCB, also leads to increased protein–water vdW interactions, producing IDP ensembles better in agreement with experimental data than the original water model.^28^ However, the detailed comparison between a99SB-ILDN/TIP4P-D and a99SB-UCB shows that the better approach is the adjustment of the LJ parameters on atom type basis with the aim to reproduce experimental solvation free energies of a diverse set of molecules^34^ instead of uniformly scaling the vdW interactions between protein and water.^28^ This conclusion is further supported by the not convincing performance of a99SB-disp, for which also the protein–water vdW interactions were increased in an amino-acid unspecific fashion.^32^ In addition, the ϕand ψparameters of all amino acids except glycine and proline were modified during the development of a99SB-disp, which together with the increased protein–water interactions led to the overestimation of the PPII propensity.

In summary, the recommendation is to use a99SB-UCB when simulating Aβ, a FF that was carefully optimized on atom-type basis by Head-Gordon and co-workers.^33,34^ If one wishes to further improve its performance for Aβ, special attention should be devoted to the sequence ^9^GYEVHH^14^ as here the deviation from the experimental *J* -couplings is the largest.

### 4.2 What does Aβ40 really look like?

An obvious question of course is what structures Aβ40 assumes in the MD simulations. As most of the FFs under consideration produced structural ensembles that are not in agreement with experiment, we limit the discussion of structures to a99SB-UCB as the best FF and C36m as one of the better FFs serving as comparison to a99SB-UCB. Given the multitude of structures hidden in an MD trajectory, it is impossible to select one structure and claim it to be the most representative one. Moreover, the aim should be to select structures that are best in agreement with experimental findings. To reach this goal we reweighted the conformations sampled by the MD simulations with a99SB-UCB and C36m using the maximum entropy principle to reduce the discrepancies between the calculated and experimental ^3^*J*_HNH*α*_ couplings (Figure S17).^57^ We observed a reduction in *χ*^2^ from 1.94 to 0.13 for a99SB-UCB and from 3.05 to 0.51 for C36m while retaining a significant fraction of the MD frames. The 50 structures with highest and the 50 ones with lowest weights, i.e., the structures best and worst in agreement with experiment were investigated in more detail. To this end, we calculated *R*_gyr_, *R*_ee_, the distance between the C_*α*_ atoms of residues K16 and D23 as well as D23 and K28.

For a99SB-UCB, the averages of these values for the high- and low-weight structures are very similar; they are within 0.2 nm of each other, and the *R*_gyr_ and *R*_ee_ values are also close to the average values of the whole trajectory. This can also be seen from the distribution of the *R*_gyr_ values, which did not change much after reweighting the MD frames (Figure S17). This indicates that large-scale motions are not responsible for the discrepancies from the NMR data of the low-weight structures, which is confirmed by Figure 7 showing the two structures with highest and the two with lowest weights for a99SB-UCB. At first glance, the high- and low-weight structures may look highly similar since they are both generally extended and disordered. However, a closer inspection reveals certain important differences. For instance, the two low-weight structures feature a F19/F20 turn not present in the two high-weight counterparts. In addition, the N-terminal regions of both groups are different: this, we believe is significant especially since the worst performance for a99SB-UCB was obtained for ^9^GYEVHH^14^. In the two high-weight structures the N-terminal sequence up to residue K16 is rather extended, while it is more collapsed in the two low-weight structures. In one of them (Figure 7D) even a helix formed between Y10 and K16, while the overall appearance of this conformation is overly compact.

**Figure 7:**
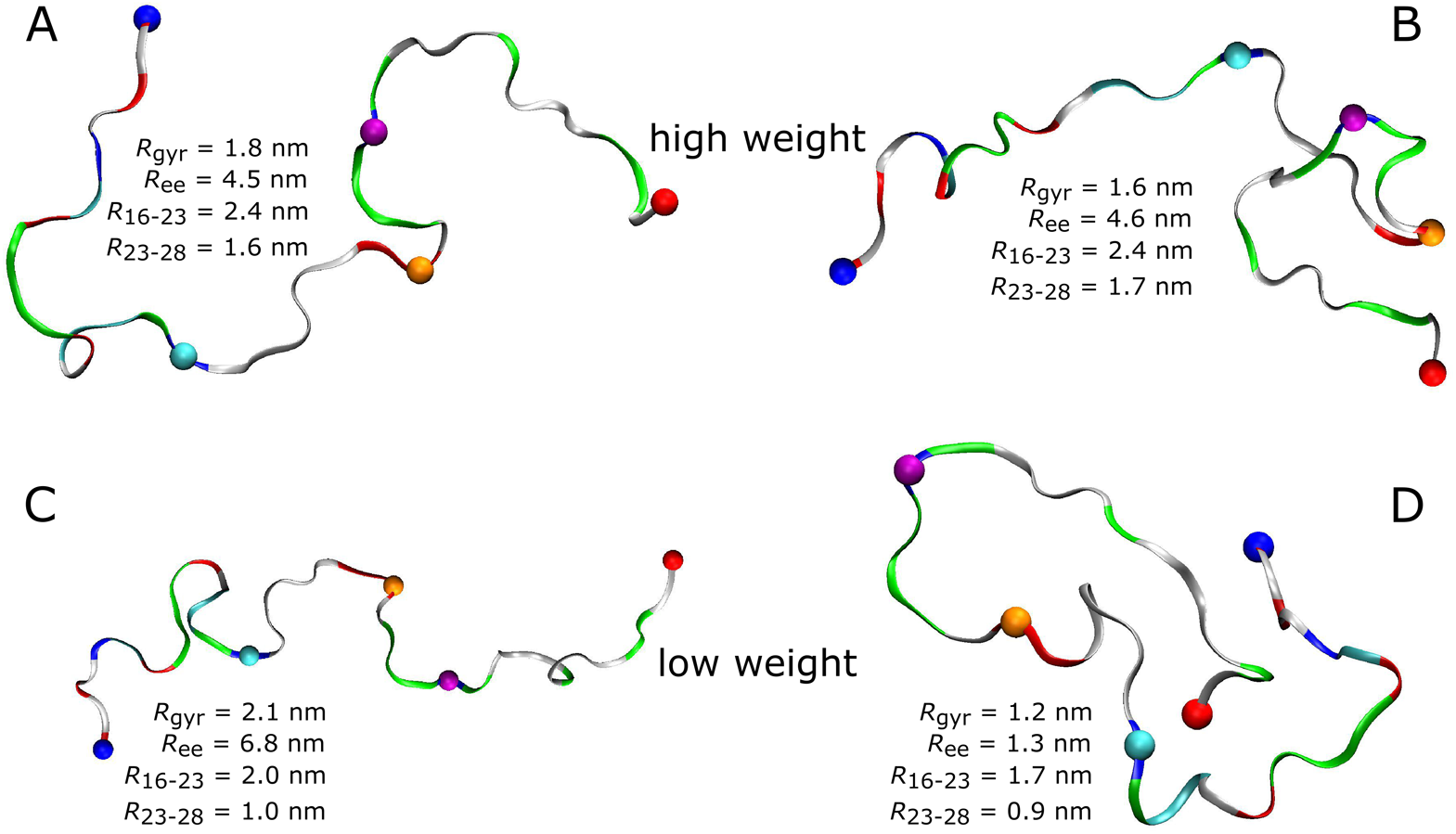
High-weight (A and B) and low-weight structures (C and D) determined by reweighting the a99SB-UCB trajectory using the Bayesian/maximum entropy technique. ^57^ Aβ40 is shown as band and colored according to amino acid residue type (basic: blue, acidic: red, histidine: cyan, polar: green, hydrophobic: white). Following residues are indicated by spheres: N-terminus (blue), K16 (cyan), D23 (orange), K28 (mauve), C-terminus (red). The structures were characterized in terms of *R*_ee_, *R*_gyr_, the K16–D23 distance (*R*_16−23_), and the D23–K28 distance (*R*_23−28_).

The inspection of the corresponding structures for C36m (Figure S18) reveals a similar structural difference for the sequence ^16^KLVFFAE^23^ as seen for a99SB-UCB. While in the high-weight structures this sequence is rather extended, it exhibits a turn at F19/F20 in the low-weight structures. Thus, based on the *J* -couplings, a turn in that region should not be sampled. Apart from this turn, visual inspection failed to identify further major differences. In fact, the conformations in Figure S18A and B (high-weight structures) and those in Figure S18C and D (low-weight structures) look rather similar. As for a99SB-UCB it is observed that reweighting the structures does not change the distribution of *R*_gyr_ values much. Nonetheless, a small change of *R*_gyr_ to a lower average value is observed for C36m. This agrees with the finding that the 50 high-weight structures obtained with C36m are on the average more compact (average *R*_ee_ = 2.79 nm) than their low-weight counterparts (average *R*_ee_ = 3.65 nm). This is accompanied by a smaller distance between D23 and K28 (0.97 nm vs. 1.24 nm) but a larger K16–D23 distance (2.11 nm vs. 1.90 nm). The conclusion is that, when a FF is not able to provide a generally satisfactory structural ensemble (as in the case of C36m), it might not be possible to optimize for different experimental observables at once; here, agreement with the *J* -couplings was optimized, which led to a larger disagreement for *R*_ee_.

If one wants to gain an in-depth understanding of the structural features sampled in the low-weight structures that are in disagreement with the various available experimental observables, it would be necessary to determine these observables for the low-weight structures (and also high-weight structures as reference) per residue and then correlate these with the structural preferences as (for instance) measured by the dihedral distribution in Ramachandran space. Based on this, suggestions for possible force field optimizations could be made. An alternative approach would be to employ an automatic reparametrization based on a Bayesian formalism that takes the available experimental data into account.^57^ However, considering that the recently attained excellent agreement between aa9SB-UCB-simulated Aβ ensemble and experimental data has taken so long to achieve by the simulation research community, it is important that extreme care is taken in performing any reoptimization attempt. The goal should be to make as little changes as necessary in order to avoid losing of what has been achieved. A classical example is shown in the case of a99SB-disp where a force field optimization procedure that was too general and/or too extensive destroyed what had already been gained in a99SB-ILDN/TIP4P-D which for Aβ40 clearly performs better than its optimized a99SB-disp descendant.

### 4.3 How long should MD simulations of Aβ be?

The kinetic analysis of the seven 30-µs MD simulation does not only allow us to conclude which of the FFs provides the best structural ensemble of Aβ, but also how long standard MD simulations of Aβ should be if one aims for converged results of the structural ensemble. To answer this question we analyzed the convergence behavior of different quantities, such as the RMSD, number of clusters, secondary structure, *R*_gyr_ and also *R*_ee_, and calculated MSMs based on TICA for dimensionality reduction. The RMSD was identified as a useless measure to assess convergence for the simulation of an IDP like Aβ. If one aims for converged results for *R*_gyr_ and *R*_ee_ it seems that for many of the FFs less than 10 µs of MD sampling is sufficient. A time limit below 10 µs has not been provided here since this was not analyzed in the present work. However, from FRET experiments of Aβ it was concluded that conformational dynamics leading to changes of the end-to-end distance does not exceed 1 µs.^35^ Thus, it might be that for this quantity a sampling time of 1 µs might even be sufficient. Yet with a03ws and a99SB-disp*-ILDN/TIP3P large changes in both *R*_ee_ and *R*_gyr_ were still observed after 10 µs of MD. However, as both FFs were already identified as not suitable for modeling the structural ensemble of Aβ, in the following discussion we will ignore them. Though it is interesting to note that FFs, which fail to provide an acceptable description of the metastable states, also fail to reproduce the peptide kinetics. This is understandable as conformations in disagreement with experiment are also expected to be connected by transition states different from reality. Moreover, the Aβ conformations favored by these two FFs are overly stable in terms of intrapeptide contacts, which explains the slow kinetics generated with them.

If one uses the secondary structure as a measure for convergence one also finds that 10 µs seem sufficient as the averages of the different structural elements did not change much beyond that time. However, the evolution of the secondary structure shows that for all FFs considerable changes in secondary structure can occur beyond 10 µs (which do not much affect the average values). Thus, the recommendation is to simulate Aβ for at least 10 µs, if possible longer. The same recommendation is made based on the cluster and MSM analyses. The number of clusters converged only beyond a sampling time of 10 µs, and for the best FF, a99SB-UCB, this was achieved only after 20 µs as a result of rare β-sheet formation, suggesting that 20 µs is a safer measure for guaranteeing that all possible conformations have been sampled. This conclusion agrees with the observations made with the MSM analysis. Here, the MFPTs suggest that 30 µs is even better as the transitions to rare states require sampling times of that length.

One might of course wonder whether 30 µs is indeed sufficient to exhaustively sample the conformational space, i.e., to consider the possibility of identifying unique conformations not sampled yet within 30 µs. We found the comparison of profiles obtained in our investigation with the MSM obtained from the 315 µs C22*/TIP3P-based MD sampling of Aβ42 very helpful in answering this question.^70^ The resulting MSM identified four macrostates with similar populations and Aβ structures as well as MFPTs as obtained from our analysis. We can thus safely conclude that there is no justifiable need to simulate monomeric Aβ for longer than 30 µs, even when kinetically slow FFs like C22*/TIP3P and C36m are being employed. The main recommendation is to use a99SB-ILDN/TIP4P-D and a99SB-UCB, and to collect 20–30 µs of MD sampling to reach conformational convergence.

## 5 Conclusions

We analyzed the 30 µs MD simulations that were acquired for Aβ40 using seven different FF/water-model combinations by D. E. Shaw Research.^32^ One aim of our analysis was to assess how much of MD time is needed for obtaining fully converged MD ensembles for this peptide. Our analysis of the evolution of different structural quantities as well as Markov state models calculated from the MD data showed that the answer to this question partly depends on the FF used as the different FFs produced different kinetics for Aβ. Since only the FFs that are in agreement with experimental results should be employed, we calculated various NMR and FRET observables from the MD trajectories and compared the resulting values to the experimental ones.^35,59^ This comparison revealed that the best FF field for Aβ is a99SB-UCB, which is based on a99SB/TIP4P-Ew and includes LJ and dihedral modifications implemented by Head-Gordon and co-workers.^33,34^ The second best performance is found for a99SB-ILDN/TIP4P-D, which can also be recommended for modeling Aβ, while all other FFs showed severe failures in reproducing at least one, in most cases more sets of experimental quantities. Usage of a03ws and a99SB*-ILDN/TIP3P for Aβ simulations is clearly discouraged as they produce too much folded Aβconformations, with too much helix in the case of a03ws and too much β-sheet with a99SB*-ILDN/TIP3P. The other three FFs under study, a99SB-disp, C22*/TIP3P, and C36m, produced acceptable results for Aβ, but considering that there are two FFs that clearly perform better, the recommendation is to use these.

The MSMs resulting from the simulations with a99SB-UCB and a99SB-ILDN/TIP4P-D are both dominated by one state that harbors extended Aβ40 structures with little to none inter-residue contacts beyond direct neighbor contacts. Two further metastable, yet low-populated states (total population of 5%) are identified with both FFs, which involve β-hairpin formation. With a99SB-UCB the β-hairpins in both these states are centered at residues V24/G25 and involve contacts between F19/F20 with I31/I32. With a99SB-ILDN/TIP4P-D such β-hairpin is also sampled in one of the MSM states, while the other one contains a β-hairpin in the N-terminal region. Transitions to these low-populated states are rare events and most of them are thus associated with MFPTs above 20 µs, reaching almost 30 µs. The connclusion therefore is that 20–30 µs of MD sampling is needed to obtain converged trajectories producing the equilibrium distribution of Aβconformations.

This conclusion is supported by the analysis of the convergence of other structural quantities, such as the number of conformational clusters sampled. However, is not necessary to sample one continuous trajectory of 20–30 µs length. Instead, several shorter trajectories could be collected, which, ideally, get initiated from different conformations. For this, the adaptive sampling scheme of MSMs could be used to identify undersampled regions from where peptide conformations can be selected for spawning new MD simulations.^71^

Unlike the MFPTs to metastable states, the implied time scales derived from an MSM are a measure for intrapeptide reorientations. For these motions, an upper limit of 1 µwas predicted from different fluorescent spectroscopic measurements.^35,36^ Only for a99SB-UCB and a99SB-ILDN/TIP4P-D the slowest implied time scales are below 1 µ, while with all other FFs the kinetics of Aβ40 is predicted to be much slower, in the cases of a03ws, a99SB-disp and a99SB*-ILDN/TIP3P the slowest intrapeptide motions even reach time scales beyond 10 µs. Thus, also in terms of kinetics a99SB-UCB and a99SB-ILDN/TIP4P-D are the only two FFs in agreement with experiment. Nonetheless, even though the thermodynamics and kinetics of Aβis modelled well with these two FFs, there is still further room for improvement. For instance, while a99SB-UCB is very good in reproducing NMR values for the C-terminal side of the peptide, this is less so the case for the region G9–H13. Thus, further FF improvements for Aβ should focus on this region, while keeping at the level of quality for the rest of the peptide. Overall, a major step forward in terms of FF quality for Aβ has been reached with a99sb-UCB and also a99SB-ILDN/TIP4P-D. It will be interesting to see how the kinetics of Aβ oligomer formation and the resulting structures will look like when simulated with these FFs, i.e., when the aggregation process is initiated from extended, disordered, and not partly folded states as was the case due to FF bias in previous Aβ aggregation simulations.

## Acknowledgement

The authors thank Ushnish Sengupta for fruitful discussions and Dr. Olujide Olubiyi for proofreading the manuscript. Computing resources granted by RWTH Aachen University under project rwth0400 is gratefully acknowledged. The funders had no role in study design, data collection and analysis, decision to publish, or preparation of the manuscript.

## Supporting Information Available

Figures showing the *R*_gyr_ distribution, the time-averaged secondary structures, sample densities for simulations extended to 35 µs, contact maps for the metastable states identified for the different force fields, NMR chemical shifts and their deviation from experimental and random coil values, Ramachandran plots of I31 and I32 obtained with the different force fields, evolution of the end-to-end distance, compact Aβ40 conformations sampled with a03ws, distribution of the FRET efficiency, implied time scales derived from the Markov state models, results from reweighting the MD frames of the a99SB-UCB and C36m trajectories using the maximum entropy principle, Aβ40 conformations sampled with C36m in agreement and disagreement with NMR *J* -coupling data.

This material is available free of charge via the Internet at http://pubs.acs.org/.

## References

(1) Dementia statistics. 2015 (accessed June 11, 2020).

(2) Oxford, A.; Stewart, E.; Rohn, T. Int. J. Alzheimers. Dis. 2020, 2020, 1–13.

(3) Burke, M. Chem. World 2014, 7253.

(4) Barage, S. H.; Sonawane, K. D. Neuropeptides 2015, 52, 1 – 18.

(5) Chiti, F.; Dobson, C. M. Annu. Rev. Biochem. 2017, 86, 27–68.

(6) Owen, M. C.; Gnutt, D.; Gao, M.; Wärmländer, S. K. T. S.; Jarvet, J.; Gräslund, A.; Winter, R.; Ebbinghaus, S.; Strodel, B. Chem. Soc. Rev. 2019, 48, 3946–3996.

(7) Marsden, I.; Minamide, L.; Bamburg, J. J. Alzheimers. Dis. 2011, 24, 681–91.

(8) Nasica-Labouze, J. et al. Chem. Rev. 2015, 115, 3518–3563.

(9) Ahmad, B.; Chen, Y.; Lapidus, L. J. Proc. Natl. Acad. Sci. U.S.A. 2012, 109, 2336–2341.

(10) Lapidus, L. J. Mol. BioSyst. 2013, 9, 29–35.

(11) Xu, Y.; Shen, J.; Luo, X.; Zhu, W.; Chen, K.; Ma, J.; Jiang, H. Proc. Natl. Acad. Sci. U.S.A. 2005, 102, 5403–5407.

(12) Olubiyi, O. O.; Strodel, B. J. Phys. Chem. B 2012, 116, 3280–3291.

(13) Sgourakis, N. G.; Yan, Y.; McCallum, S. A.; Wang, C.; García, A. E. J. Mol. Biol. 2007, 368, 1448–1457.

(14) Sgourakis, N. G.; Merced-Serrano, M.; Boutsidis, C.; Drineas, P.; Du, Z.; Wang, C.; García, A. E. J. Mol. Biol. 2011, 405, 570–583.

(15) Somavarapu, A. K.; Kepp, K. P. ChemPhysChem 2015, 16, 3278–3289.

(16) Gerben, S. R.; Lemkul, J. A.; Brown, A. M.; Bevan, D. R. J. Biomol. Struct. Dyn. 2014, 32, 1817–1832.

(17) Carballo-Pacheco, M.; Strodel, B. Protein Sci. 2017, 26, 174–185.

(18) Jorgensen, W. L.; Tirado-Rives, J. J. Am. Chem. Soc. 1988, 110, 1657–1666.

(19) Kaminski, G. A.; Friesner, R. A.; Tirado-Rives, J.; Jorgensen, W. L. J. Phys. Chem. B 2001, 105, 6474–6487.

(20) Jorgensen, W. L.; Chandrasekhar, J.; Madura, J. D.; Impey, R. W.; Klein, M. L. J. Chem. Phys. 1983, 79, 926–935.

(21) Hornak, V.; Abel, R.; Okur, A.; Strockbine, B.; Roitberg, A.; Simmerling, C. PROTEINS 2006, 65, 712–725.

(22) Horn, H. W.; Swope, W. C.; Pitera, J. W.; Madura, J. D.; Dick, T. J.; Hura, G. L.; Head-Gordon, T. J. Chem. Phys. 2004, 120, 9665–9678.

(23) Best, R. B.; Hummer, G. J. Phys. Chem. B 2009, 113, 9004–9015.

(24) Lindorff-Larsen, K.; Piana, S.; Palmo, K.; Maragakis, P.; Klepeis, J. L.; Dror, R. O.; Shaw, D. E. PROTEINS 2010, 78, 1950–1958.

(25) Li, D.-W.; Brüschweiler, R. Angew. Chem., Int. Ed. 2010, 49, 6778–6780.

(26) Piana, S.; Lindorff-Larsen, K.; Shaw, D. E. Biophys. J. 2011, 100, L47–L49.

(27) Rauscher, S.; Gapsys, V.; Gajda, M. J.; Zweckstetter, M.; de Groot, B. L.; Grubmüller, H. J. Chem. Theory Comput. 2015, 11, 5513–5524.

(28) Piana, S.; Donchev, A. G.; Robustelli, P.; Shaw, D. E. J. Phys. Chem. B 2015, 119, 5113–5123.

(29) Wang, W.; Ye, W.; Jiang, C.; Luo, R.; Chen, H.-F. Chem. Biol. Drug Des. 2014, 84, 253–269.

(30) Best, R. B.; Zheng, W.; Mittal, J. J. Chem. Theory Comput. 2014, 10, 5113–5124.

(31) Huang, J.; Rauscher, S.; Nawrocki, G.; Ting, R.; Feig, M.; de Groot, B.; Grubmüller, H.; MacKerell, A. Biophys. J. 2017, 112.

(32) Robustelli, P.; Piana, S.; Shaw, D. Proc. Natl. Acad. Sci. U.S.A. 2018, 115, 201800690.

(33) Nerenberg, P. S.; Jo, B.; So, C.; Tripathy, A.; Head-Gordon, T. J. Phys. Chem. B 2012, 116, 4524–4534.

(34) Nerenberg, P. S.; Head-Gordon, T. J. Chem. Theory Comput. 2011, 7, 1220–1230.

(35) Meng, F.; Bellaiche, M. M.; Kim, J. Y.; Zerze, G. H.; Best, R. B.; Chung, H. S. Biophys. J. 2018,

(36) Acharya, S.; Srivastava, K.; Nagarajan, S.; Lapidus, L. ChemPhysChem 2016, 17, 3470–3479.

(37) Watson, A. A.; Fairlie, D. P.; Craik, D. J. Biochemistry 1998, 37, 12700–12706.

(38) Van Der Spoel, D.; Lindahl, E.; Hess, B.; Groenhof, G.; Mark, A. E.; Berendsen, H. J. GROMACS: Fast, flexible, and free. 2005.

(39) Pronk, S.; Páll, S.; Schulz, R.; Larsson, P.; Bjelkmar, P.; Apostolov, R.; Shirts, M. R.; Smith, J. C.; Kasson, P. M.; Van Der Spoel, D.; Hess, B.; Lindahl, E. Bioinformatics 2013,

(40) Humphrey, W.; Dalke, A.; Schulten, K. J. Mol. Graph. 1996,

(41) Michaud-Agrawal, N.; Denning, E. J.; Woolf, T. B.; Beckstein, O. J. Comput. Chem. 2011,

(42) McGibbon, R. T.; Beauchamp, K. A.; Harrigan, M. P.; Klein, C.; Swails, J. M.; Hernández, C. X.; Schwantes, C. R.; Wang, L. P.; Lane, T. J.; Pande, V. S. Biophys. J. 2015,

(43) Bowers, K. J.; Chow, E.; Xu, H.; Dror, R. O.; Eastwood, M. P.; Gregersen, B. A.; Klepeis, J. L.; Kolossvary, I.; Moraes, M. A.; Sacerdoti, F. D.; Salmon, J. K.; Shan, Y.; Shaw, D. E. Scalable algorithms for molecular dynamics simulations on commodity clusters. Proc. 2006 ACM/IEEE Conf. Supercomput. SC’06. 2006.

(44) Daura, X.; Gademann, K.; Jaun, B.; Seebach, D.; van Gunsteren, W. F.; Mark, A. E. Angew. Chem., Int. Ed. 1999,

(45) Frishman, D.; Argos, P. PROTEINS 1995,

(46) Scherer, M. K.; Trendelkamp-Schroer, B.; Paul, F.; Pérez-Hernández, G.; Hoffmann, M.; Plattner, N.; Wehmeyer, C.; Prinz, J.-H.; Noé, F. J. Chem. Theory Comput. 2015, 11, 5525–5542.

(47) Scherer, M. K.; Husic, B. E.; Hoffmann, M.; Paul, F.; Wu, H.; Noé, F. J. Chem. Phys. 2019, 150, 194108.

(48) Pérez-Hernández, G.; Paul, F.; Giorgino, T.; De Fabritiis, G.; Noé, F. J. Chem. Phys. 2013, 139, 015102.

(49) Campello, R.; Moulavi, D.; Sander, J. Density-Based Clustering Based on Hierarchical Density Estimates. Advances in Knowledge Discovery and Data Mining. PAKDD 2013. Lecture Notes in Computer Science. 2013; pp 160–172.

(50) Kube, S.; Weber, M. J. Chem. Phys. 2007, 126, 024103.

(51) Röblitz, S.; Weber, M. Adv. Data Anal. Classi. 2013, 7, 147–179.

(52) Prinz, J.-H.; Wu, H.; Sarich, M.; Keller, B.; Senne, M.; Held, M.; Chodera, J. D.; Schütte, C.; Noé, F. J. Chem. Phys. 2011, 134, 174105.

(53) Shen, Y.; Bax, A. J. Biomol. NMR 2010,

(54) Karplus, M. J. Chem. Phys. 1959,

(55) Vögeli, B.; Ying, J.; Grishaev, A.; Bax, A. J. Am. Chem. Soc. 2007,

(56) Lincoff, J.; Sasmal, S.; Head-Gordon, T. J. Chem. Phys. 2019, 150, 104108.

(57) Bottaro, S.; Bengtsen, T.; Lindorff-Larsen, K. In Structural Bioinformatics: Methods and Protocols; Gáspári, Z., Ed.; Springer US: New York, NY, 2020; pp 219–240.

(58) Larsen, A. H.; Wang, Y.; Bottaro, S.; Grudinin, S.; Arleth, L.; Lindorff-Larsen, K. PLoS Comput. Biol. 2020, 16, 1–29.

(59) Roche, J.; Shen, Y.; Lee, J. H.; Ying, J.; Bax, A. Biochemistry 2016, 55, 762–775.

(60) Wishart, D.; Case, D. Methods Enzymol. 2001, 338, 3–34.

(61) Wang, Y.; Jardetzky, O. J. Am. Chem. Soc. 2002, 124, 14075–14084.

(62) Mantsyzov, A. B.; Maltsev, A. S.; Ying, J.; Shen, Y.; Hummer, G.; Bax, A. Protein Sci. 2014, 23, 1275–1290.

(63) Mantsyzov, A.; Shen, Y.; Lee, J.; Hummer, G.; Bax, A. J. Biomol. NMR 2015, 63, 85–95.

(64) Kjaergaard, M.; Poulsen, F. J. Biomol. NMR 2011, 50, 157–65.

(65) Kjaergaard, M.; Brander, S.; Poulsen, F. J. Biomol. NMR 2011, 49, 139–49.

(66) Best, R.; Buchete, N.-V.; Hummer, G. Biophys. J. 2008, 95, L07–9.

(67) Granata, D.; Baftizadeh, F.; Habchi, J.; Galvagnion, C.; De Simone, A.; Camilloni, C.; Laio, A.; Vendruscolo, M. Sci. Rep. 2015, 5, 15449.

(68) Nygaard, M.; Kragelund, B. B.; Papaleo, E.; Lindorff-Larsen, K. Biophys. J. 2017, 113, 550 – 557.

(69) Rezaei-Ghaleh, N.; Parigi, G.; Zweckstetter, M. J. Phys. Chem. Lett. 2019, 10, 3369–3375.

(70) Löhr, T.; Kohlhoff, K.; Heller, G.; Camilloni, C.; Vendruscolo, M. bioRxiv 2020,

(71) Bowman, G. R.; Ensign, D. L.; Pande, V. S. J. Chem. Theory Comput. 2010, 6, 787–794.

