## Supporting Information for "Thermodynamics and kinetics of the amyloid-β peptide revealed by Markov state models based on MD data in agreement with experiment"

July 27, 2020

### Supplementary Figures


S1 Distribution of the radius of gyration $R_{\mathrm{gyr}}$ for increasing trajectory lengths (0–10 µs: yellow, 0–20 µs: cyan, 0–30 µs: magenta) for the different force fields (labels on the top of the panels). . . . . . . . . . . . . . . . . . . . . . . . . . S4

S2 Time-averaged secondary structures coil, turn, β-sheet, and α-helix for increasing trajectory lengths (0–10 µs: yellow, 0–20 µs: cyan, 0–30 µs: magenta) for the different force fields (labels on the top of the panels). . . . . . . . . . . . . . . . S5

S3 Sample densities for different time windows of the trajectories (0–10 µs: yellow, 10–20 µs: cyan, 20–30 µs: magenta, 30–35 µs: green) projected along the first two TICA components for a99SB-disp (left) and C36m (right). . . . . . . . . . . S6

S4 Normalized contacts for the coarse-grained MSM states obtained from the simulation with a03ws. . . . . . . . . . . . . . . . . . . . . . . . . . . . . . S7

S5 Normalized contacts for the coarse-grained MSM states obtained from the simulation with a99SB-ILDN/TIP4P-D. . . . . . . . . . . . . . . . . . . . . . . . . S8

S6 Normalized contacts for the coarse-grained MSM states obtained from the simulation with a99SB-UCB. . . . . . . . . . . . . . . . . . . . . . . . . . . . S9

S7 Normalized contacts for the coarse-grained MSM states obtained from the simulation with a99SB-disp. . . . . . . . . . . . . . . . . . . . . . . . . . . . S10

S8 Normalized contacts for the coarse-grained MSM states obtained from the simulation with C22*/TIP3P. . . . . . . . . . . . . . . . . . . . . . . . . . . . S11

S9 Normalized contacts for the coarse-grained MSM states obtained from the simulation with C36m. . . . . . . . . . . . . . . . . . . . . . . . . . . . . . . S12

S10 Experimental (black) and calculated (colored) NMR chemical shifts for the C′ atoms (left) and $C_\alpha$ atoms (right) for the different force fields (indicated on the left of each row). . . . . . . . . . . . . . . . . . . . . . . . . . . . . . . . . S13

S11 Difference between (A) the calculated and experimental and (B) the calculated and random coil NMR chemical shifts for the C′ atoms of Aβ40 residues for the different force fields (color key on the bottom). The residues with the largest deviation between simulation and experiment are labeled in panel A. . . . . . . . S14

S12 Ramachandran plots of I31 and I32 obtained from the simulation with a03ws (top) and a99SB-ILDN/TIP4P-D (bottom). . . . . . . . . . . . . . . . . . . . . S15

S12 (cont.) Ramachandran plots of I31 and I32 obtained from the simulation with a99SB-UCB (top) and a99SB-disp (bottom). . . . . . . . . . . . . . . . . . . . S16

S12 (cont.) Ramachandran plots of I31 and I32 obtained from the simulation with a99SB*-ILDN/TIP3P (top) and C22*/TIP3P (bottom). . . . . . . . . . . . . . S17

S12 (cont.) Ramachandran plots of I31 and I32 obtained from the simulation with C36m. . . . . . . . . . . . . . . . . . . . . . . . . . . . . . . . . . . . . . S18

S13 Evolution the end-to-end distance $R_{\mathrm{ee}}$ for the different force fields (labels on the top of the panels). . . . . . . . . . . . . . . . . . . . . . . . . . . . . . S19

S14 Compact Aβ40 structures sampled with a03ws between 16 and 25 μs. These conformations exhibit a high propenisty for helix formation in different parts along the sequence: (A) between residues K16 and K28 (as present in MSM state 1), (B) between residues G29 and M35 (as present in MSM states 3), (C) between residues K16 to K28 and G29 to M35 (as present in MSM state 2). Aβ40 is shown as band and colored according to amino acid residue type (basic: blue, acidic: red, histidine: cyan, polar: green, hydrophobic: white). Following residues are indicated by spheres: N-terminus (blue), K16 (cyan), D23 (orange), K28 (mauve), C-terminus (red). . . . . . . . . . . . . . . . . . . . . . . . . . . S20

S15 Distribution of the FRET efficiency $E_{\mathrm{FRET}}$ for increasing trajectory lengths (0–10 μs: yellow, 0–20 μs: cyan, 0–30 μs: magenta) for the different force fields (labels on the top of the panels). . . . . . . . . . . . . . . . . . . . . . . . . . S21

S16 Implied time scales of the slowest processes (colored lines) obtained for different MSMs at different lag times (dots on colored lines) calculated from the MD trajectories using different force fields (labels on the top of the panels). . . . . . S22

S17 Reweighting of the trajectory frames using the maximum entropy principle to optimize the $J$-couplings obtained with the MD trajectory with a99SB-UCB (left) and C36m (right). (Top) The black dots indicate the experimental $J$-couplings for the individual Aβ40 residues (sorted in increasing $J$-coupling order), blue and red dots indicate the calculated $J$-couplings before and after, respectively, reweighting. (Bottom) Distribution of the radius of gyration before (blue) and after (red) reweighting the MD frames. The vertical lines indicate the corresponding $R_{\mathrm{gyr}}$ average. . . . . . . . . . . . . . . . . . . . . . . . . . . . . S23

S18 High-weight (A and B) and low-weight structures (C and D) determined by reweighting the C36m trajectory using the Bayesian/maximum entropy technique. Aβ40 is shown as band and colored according to amino acid residue type (basic: blue, acidic: red, histidine: cyan, polar: green, hydrophobic: white). Following residues are indicated by spheres: N-terminus (blue), K16 (cyan), D23 (orange), K28 (mauve), C-terminus (red). The structures were characterized in terms of $R_{\mathrm{ee}}$, $R_{\mathrm{gyr}}$, the K16–D23 distance ($R_{16-23}$), and the D23–K28 distance ($R_{23-28}$). . . . . . . . . . . . . . . . . . . . . . . . . . . . . . . . . . . S24

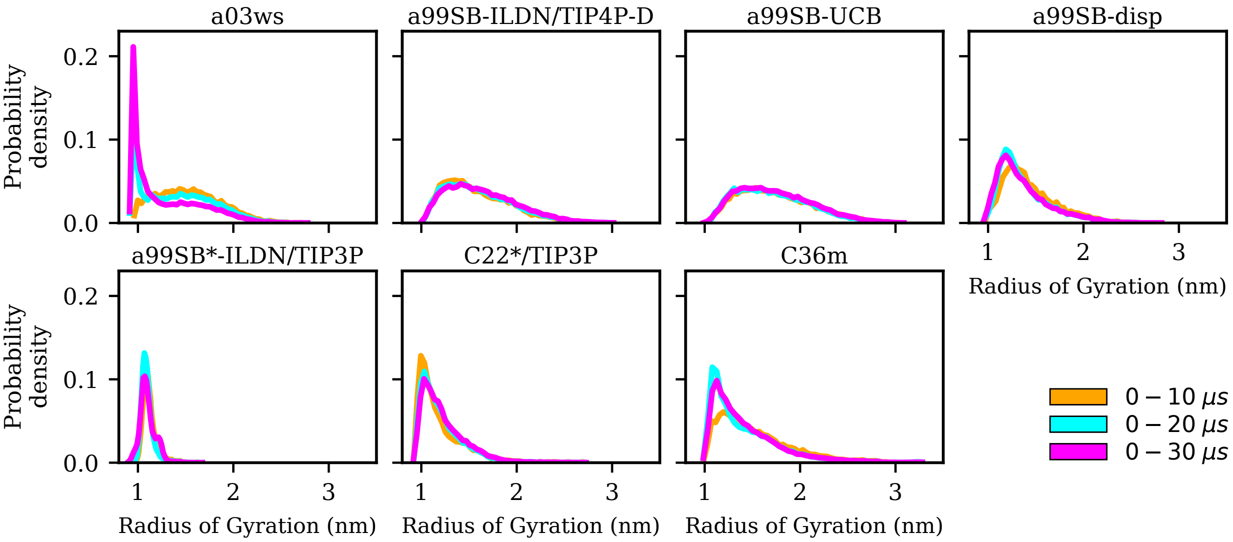


Figure S1: Distribution of the radius of gyration $R_{\text{gyr}}$ for increasing trajectory lengths (0–10 µs: yellow, 0–20 µs: cyan, 0–30 µs: magenta) for the different force fields (labels on the top of the panels).

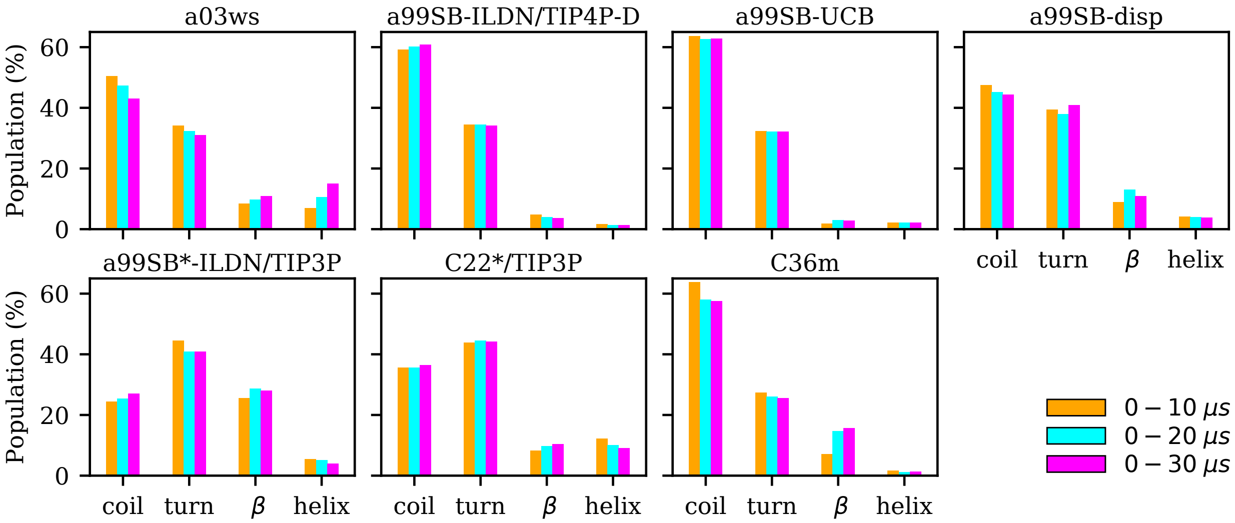


Figure S2: Time-averaged secondary structures coil, turn, β-sheet, and α-helix for increasing trajectory lengths (0–10 μs: yellow, 0–20 μs: cyan, 0–30 μs: magenta) for the different force fields (labels on the top of the panels).

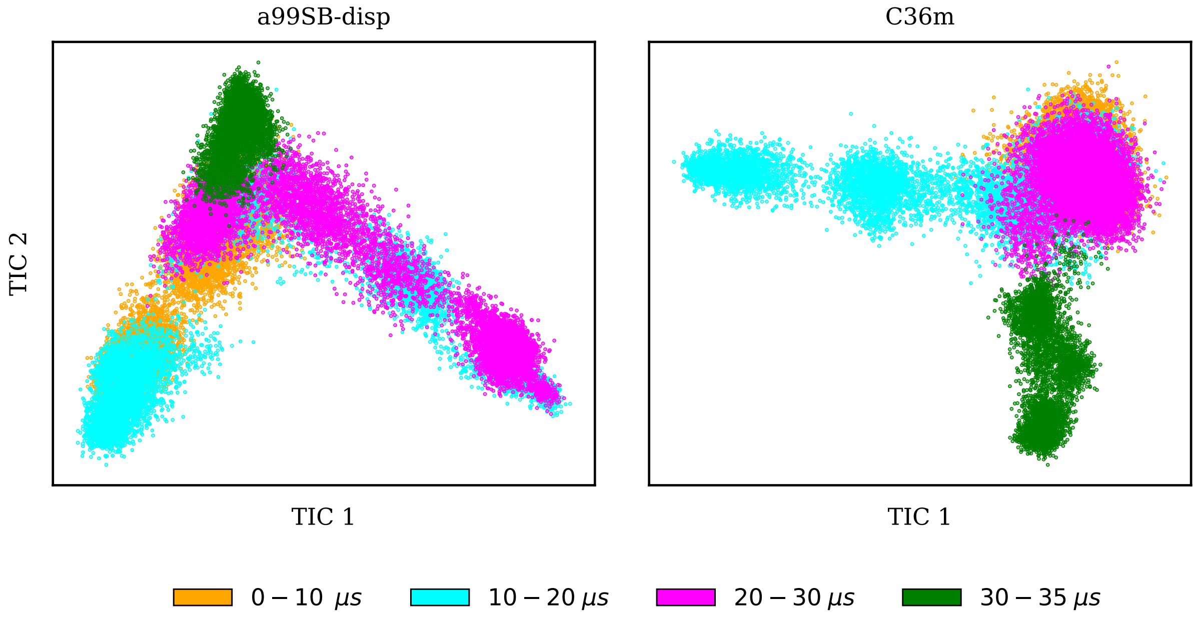


Figure S3: Sample densities for different time windows of the trajectories (0–10 μs: yellow, 10–20 μs: cyan, 20–30 μs: magenta, 30–35 μs: green) projected along the first two TICA components for a99SB-disp (left) and C36m (right).

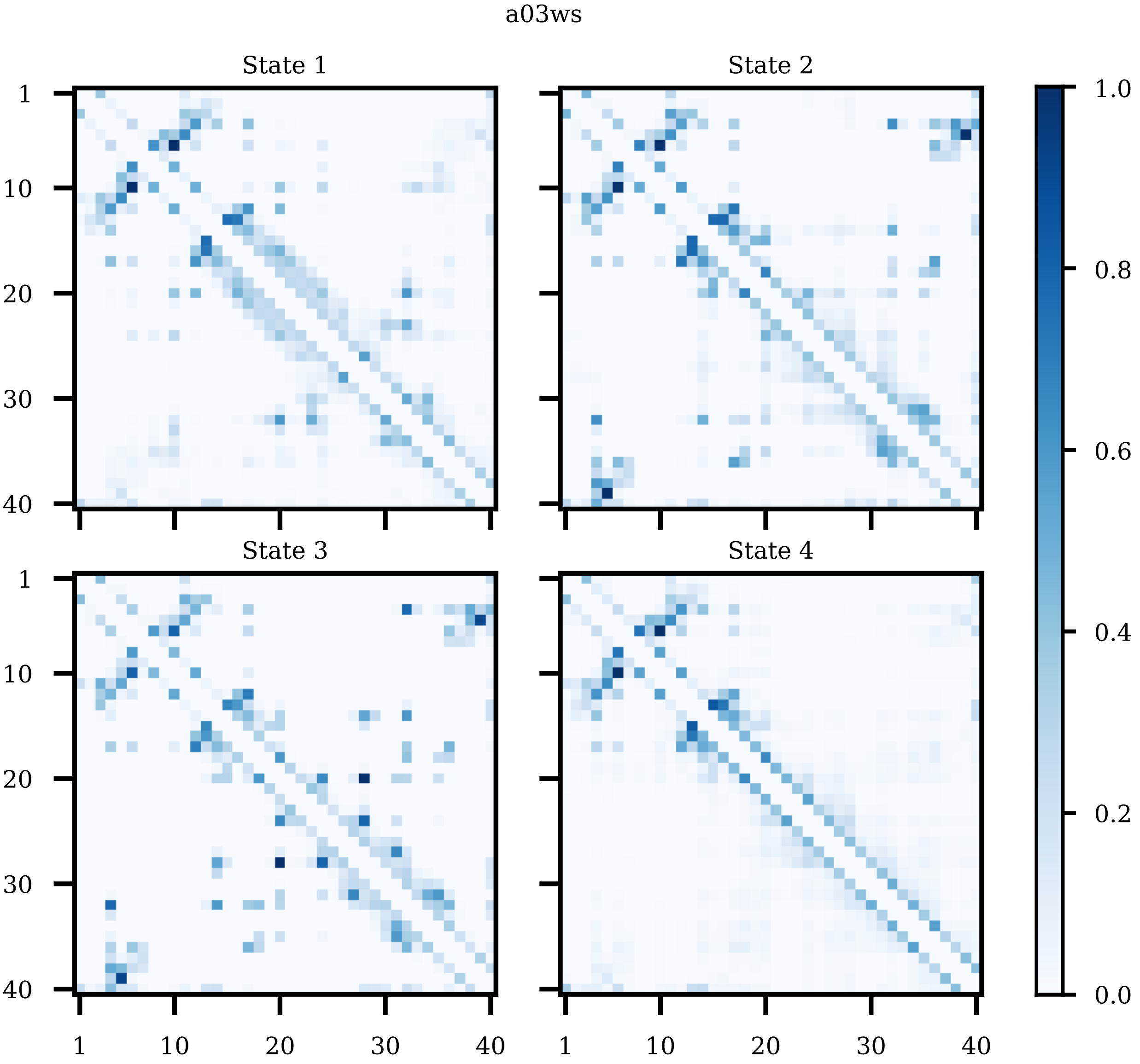


Figure S4: Normalized contacts for the coarse-grained MSM states obtained from the simulation with a03ws.

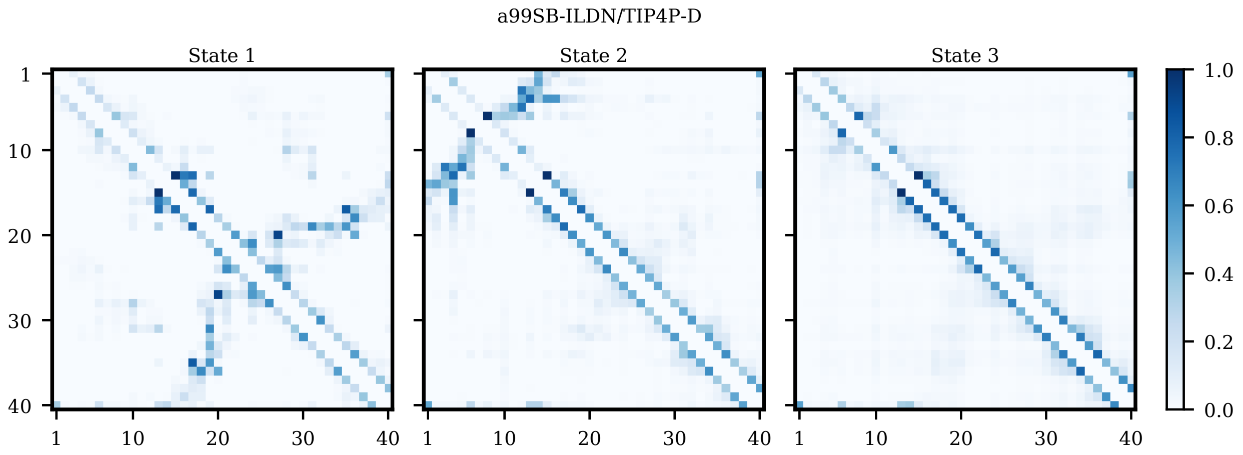


Figure S5: Normalized contacts for the coarse-grained MSM states obtained from the simulation with a99SB-ILDN/TIP4P-D.

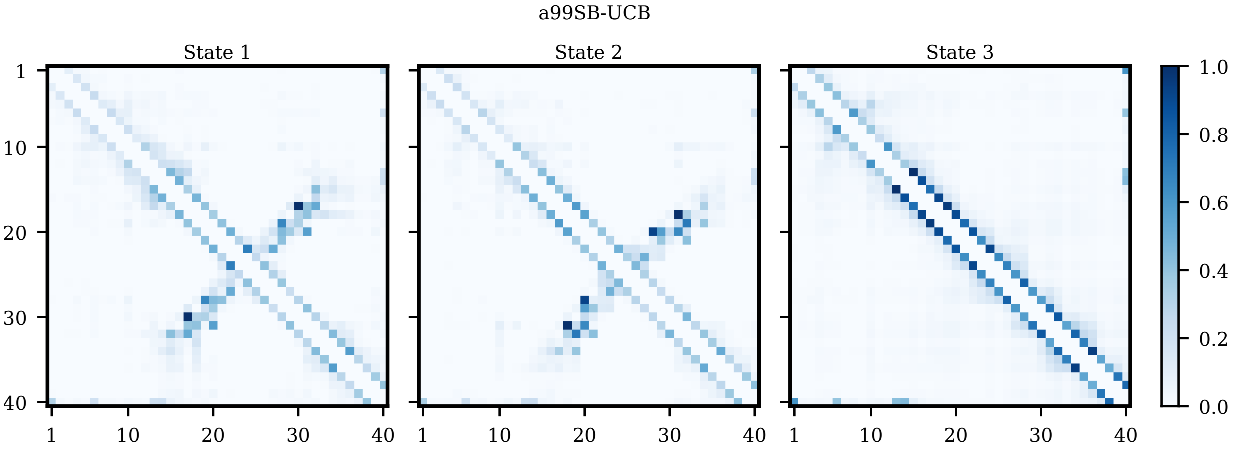


Figure S6: Normalized contacts for the coarse-grained MSM states obtained from the simulation with a99SB-UCB.

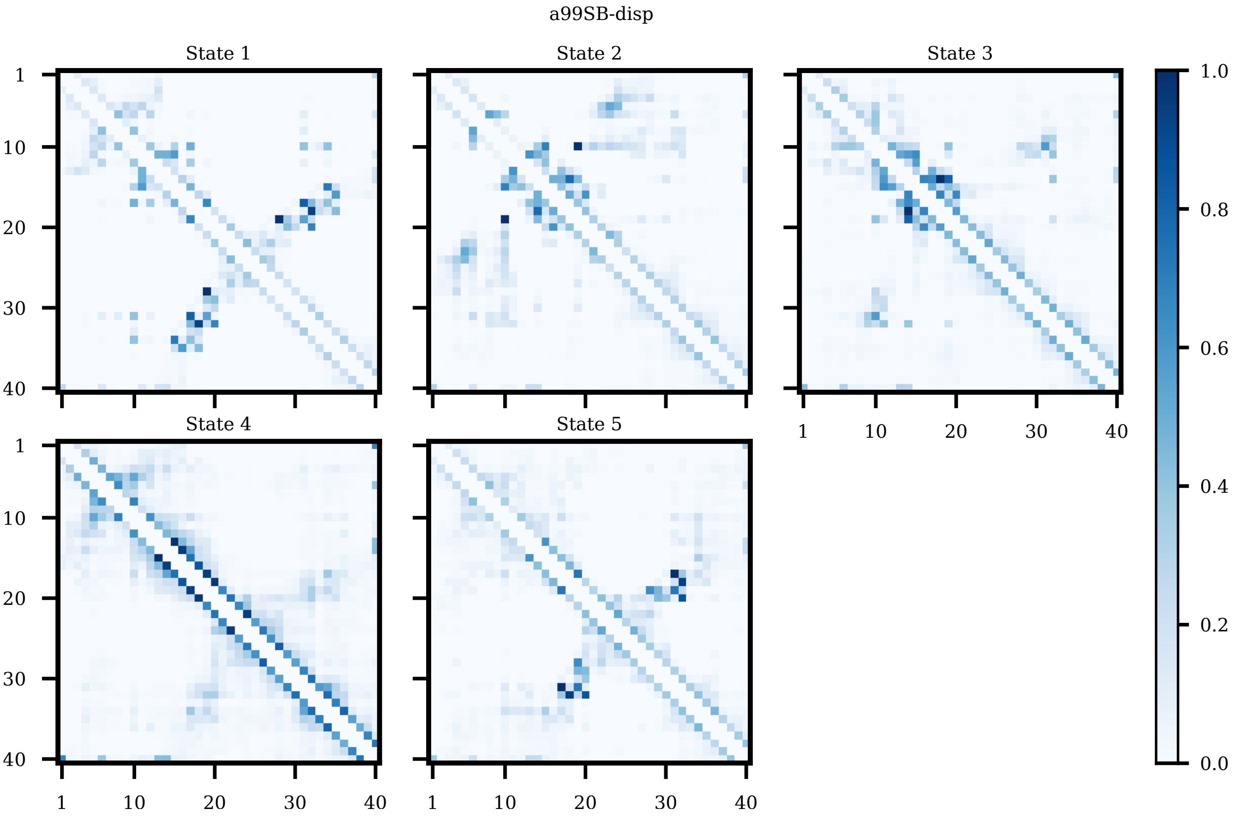


Figure S7: Normalized contacts for the coarse-grained MSM states obtained from the simulation with a99SB-disp.

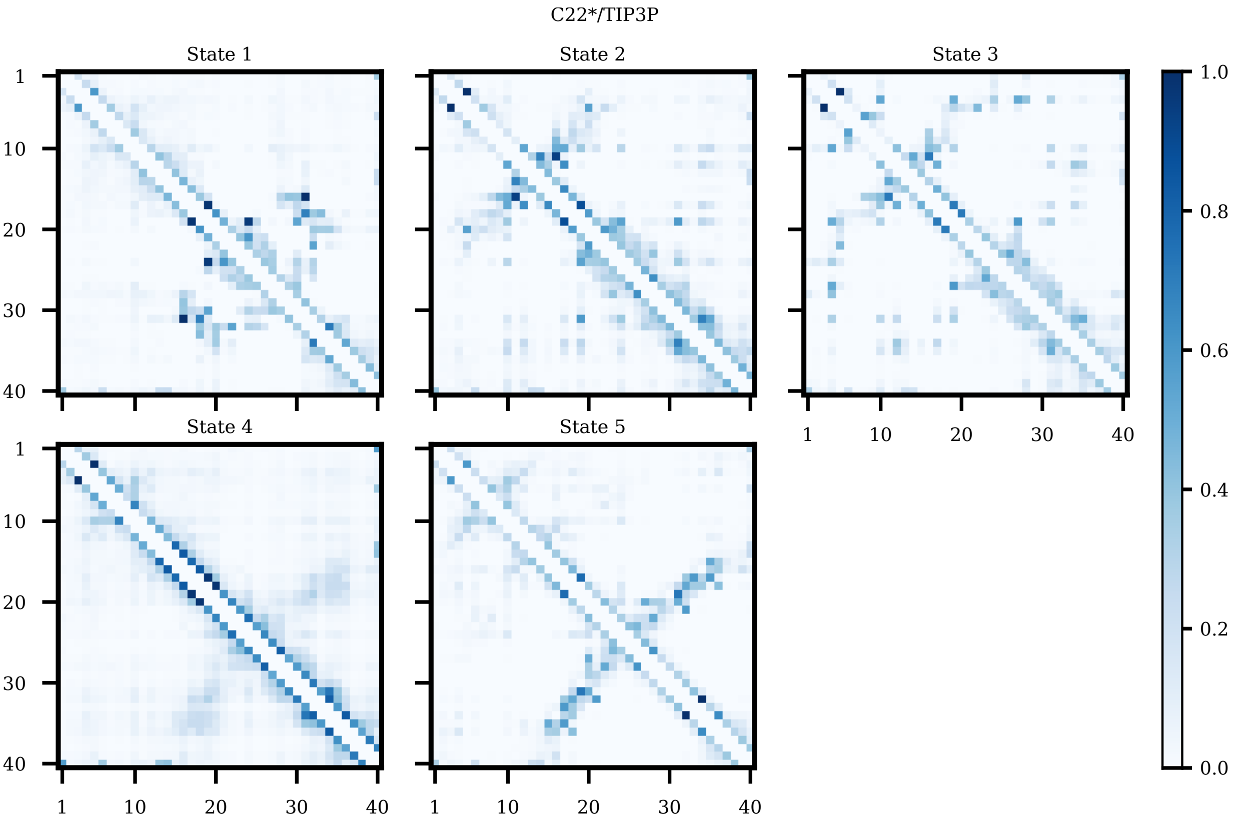


Figure S8: Normalized contacts for the coarse-grained MSM states obtained from the simulation with C22*/TIP3P.

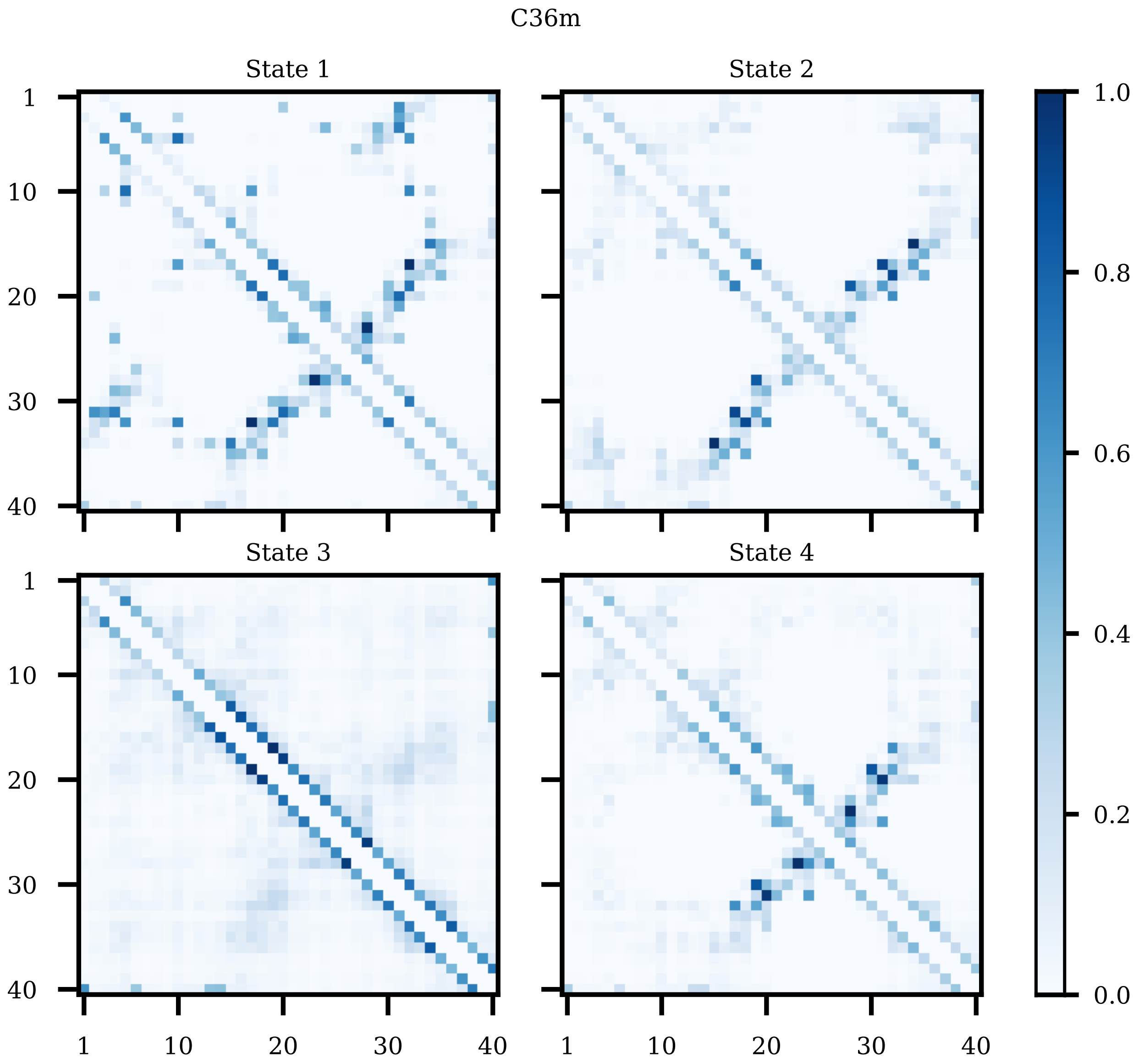


Figure S9: Normalized contacts for the coarse-grained MSM states obtained from the simulation with C36m.

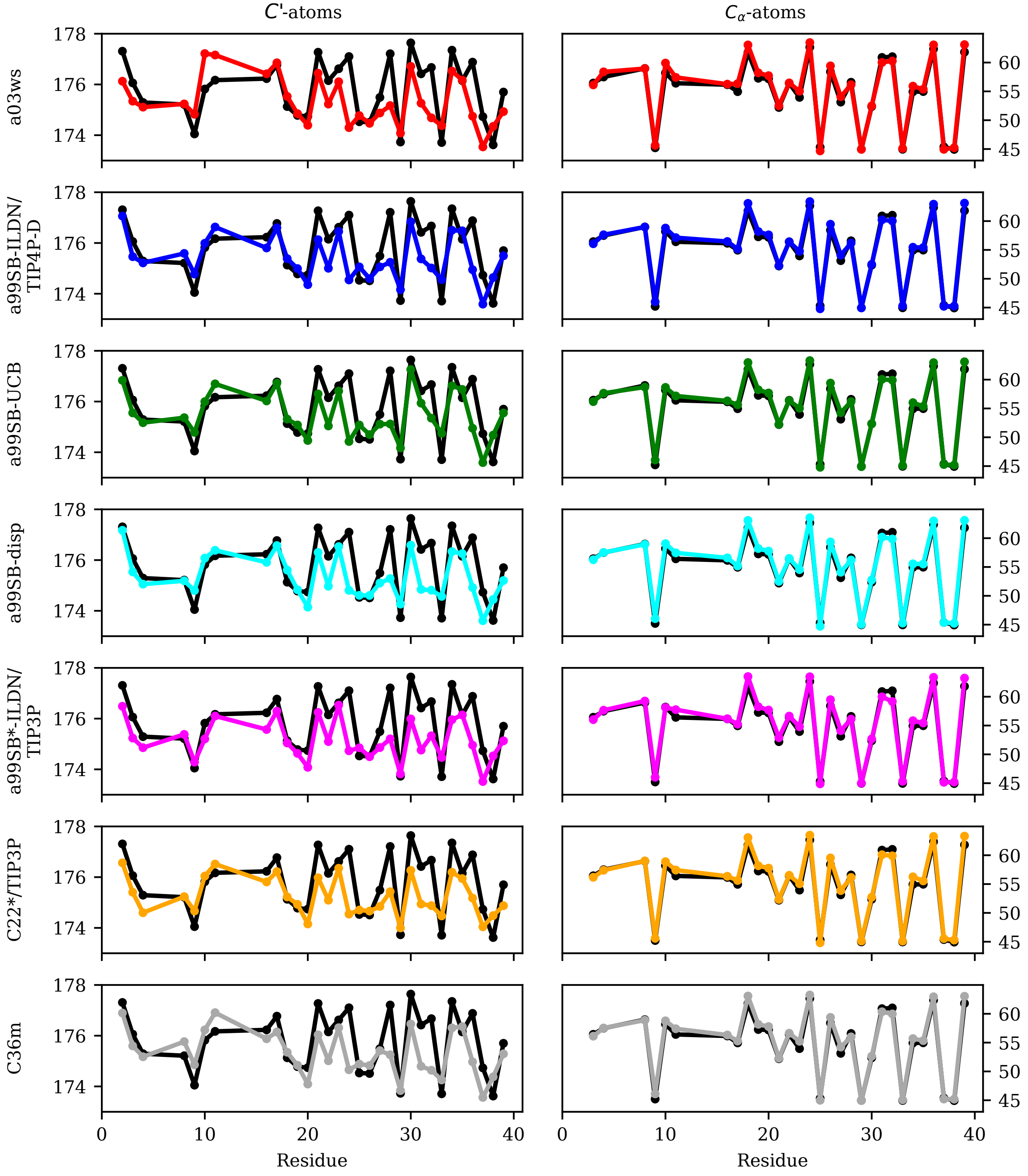


Figure S10: Experimental (black) and calculated (colored) NMR chemical shifts for the C′ atoms (left) and $C_\alpha$ atoms (right) for the different force fields (indicated on the left of each row).

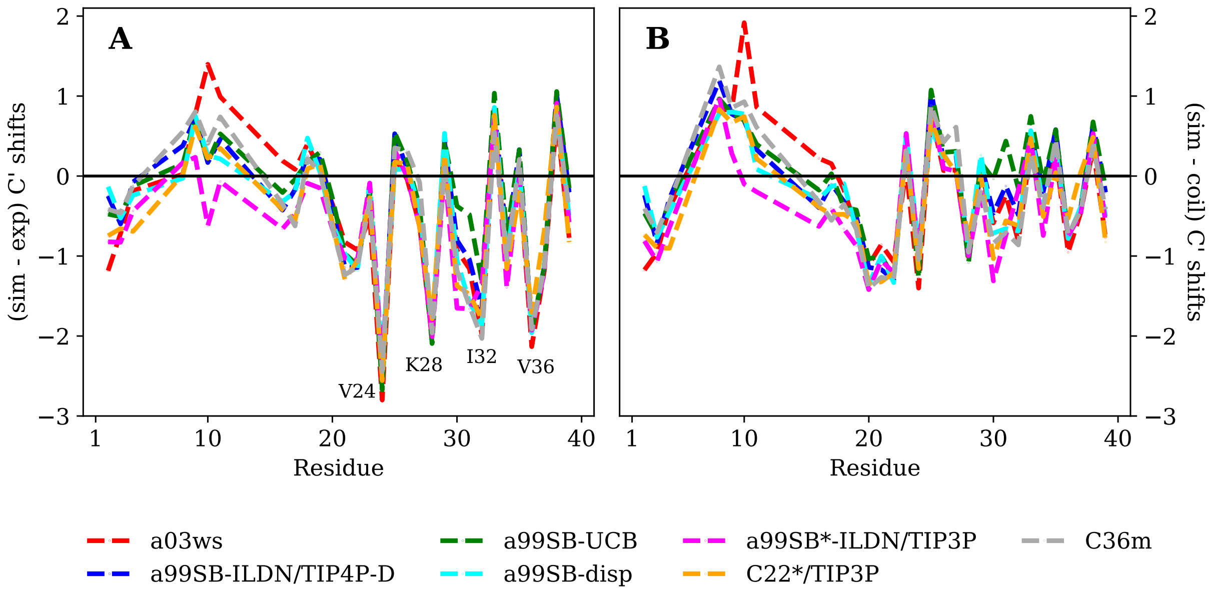


Figure S11: Difference between (A) the calculated and experimental and (B) the calculated and random coil NMR chemical shifts for the C′ atoms of Aβ40 residues for the different force fields (color key on the bottom). The residues with the largest deviation between simulation and experiment are labeled in panel A.

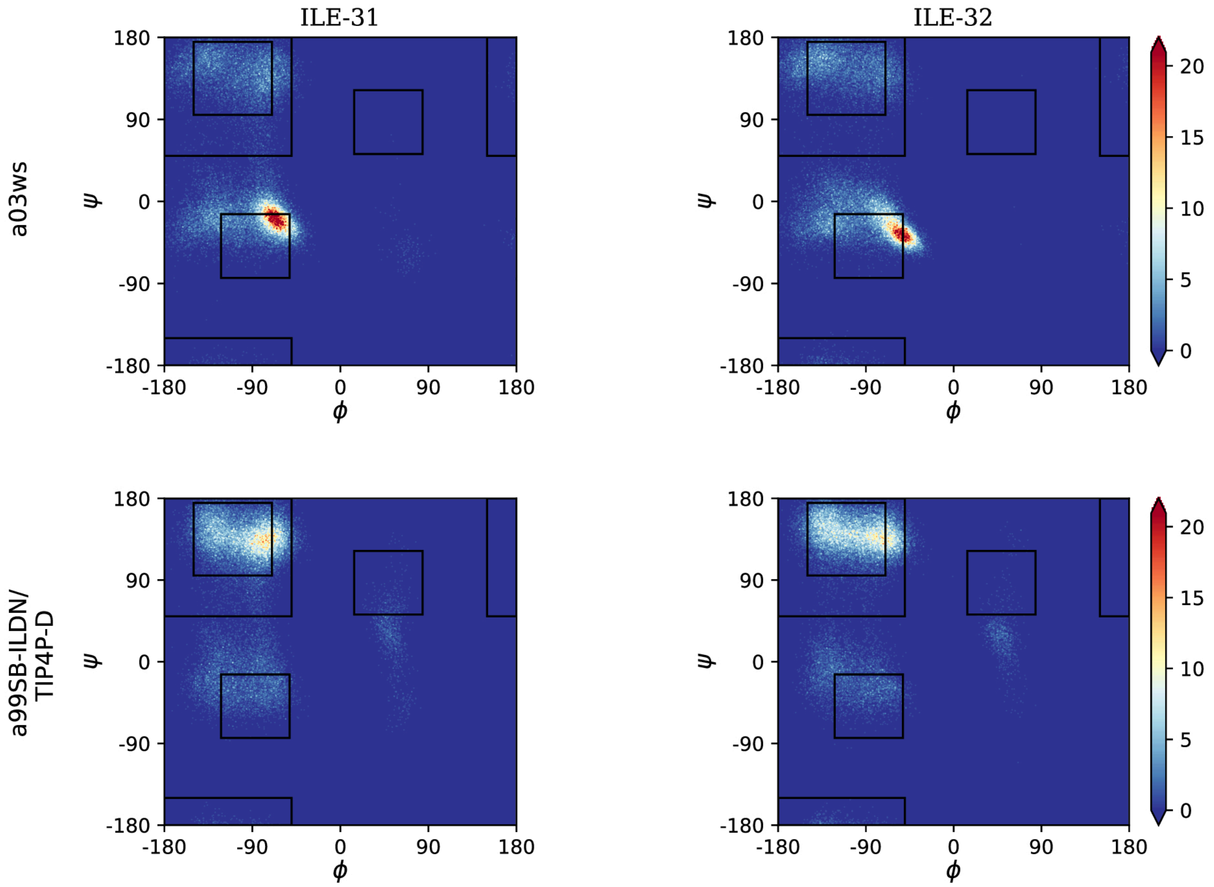


Figure S12: Ramachandran plots of I31 and I32 obtained from the simulation with a03ws (top) and a99SB-ILDN/TIP4P-D (bottom).

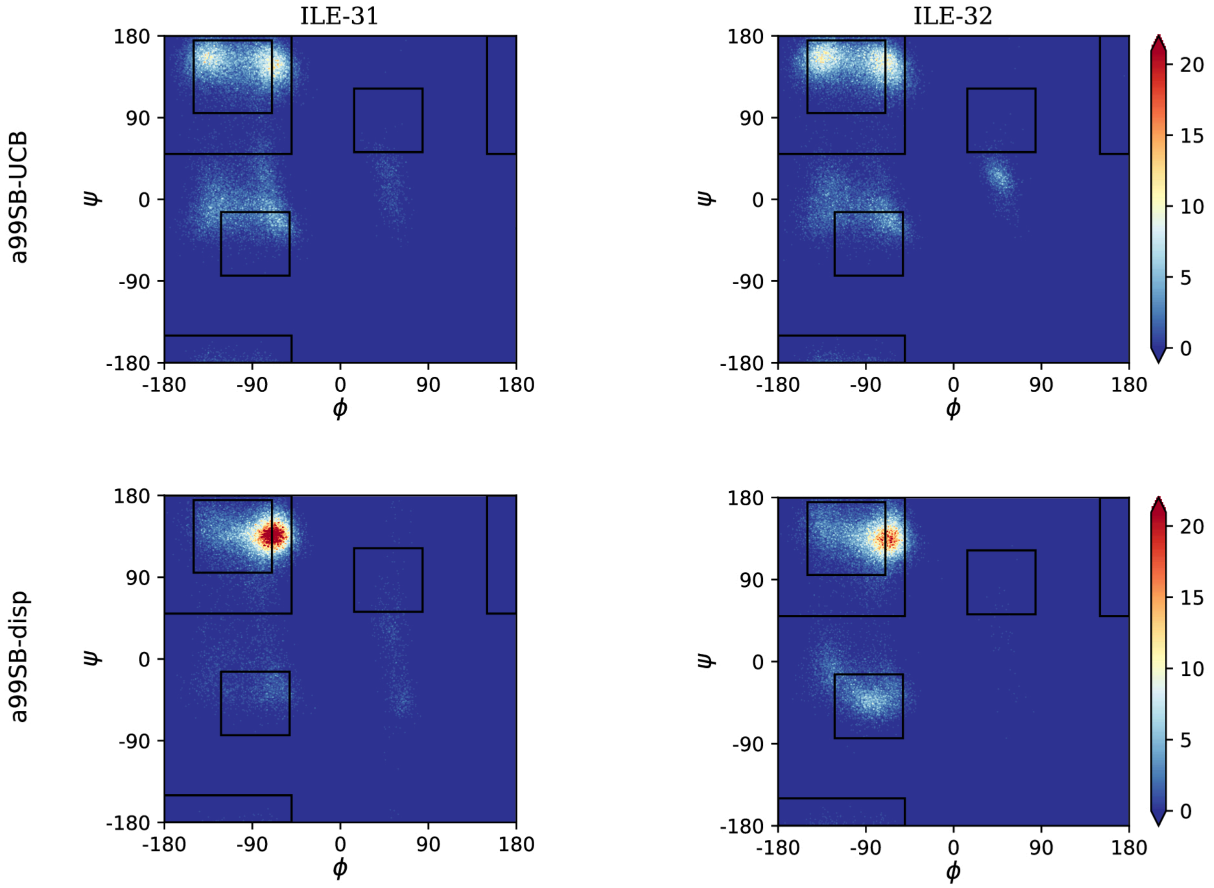


Figure S12: (cont.) Ramachandran plots of I31 and I32 obtained from the simulation with a99SB-UCB (top) and a99SB-disp (bottom).

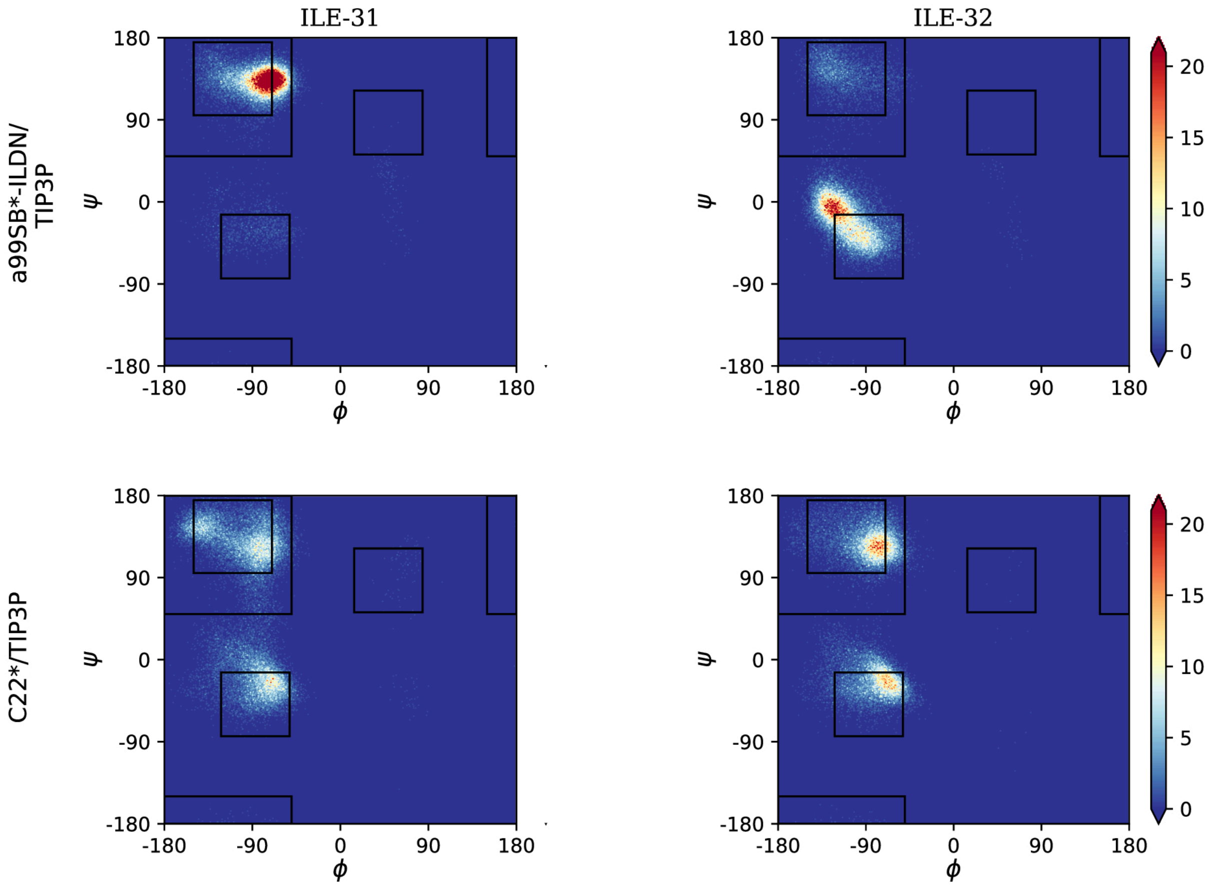


Figure S12: (cont.) Ramachandran plots of I31 and I32 obtained from the simulation with a99SB*-ILDN/TIP3P (top) and C22*/TIP3P (bottom).

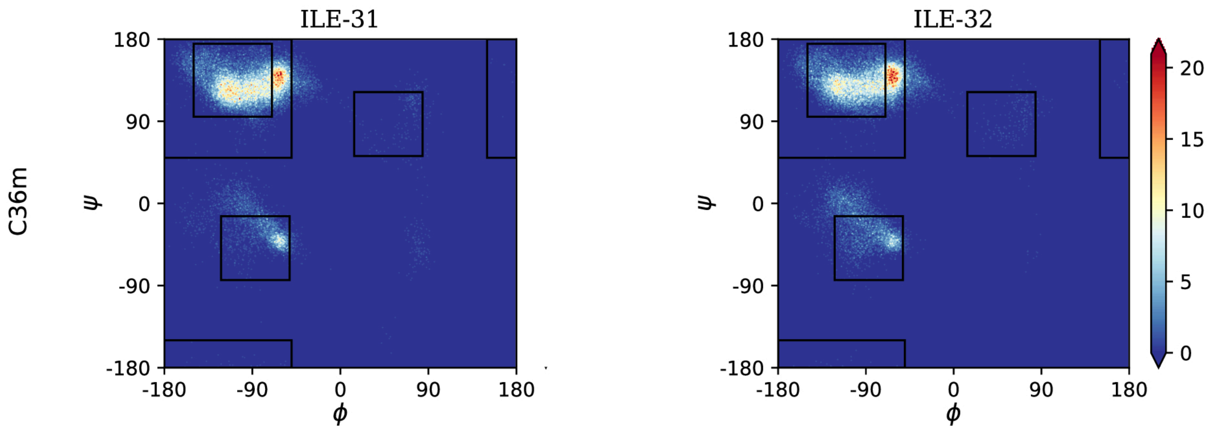


Figure S12: (cont.) Ramachandran plots of I31 and I32 obtained from the simulation with C36m.

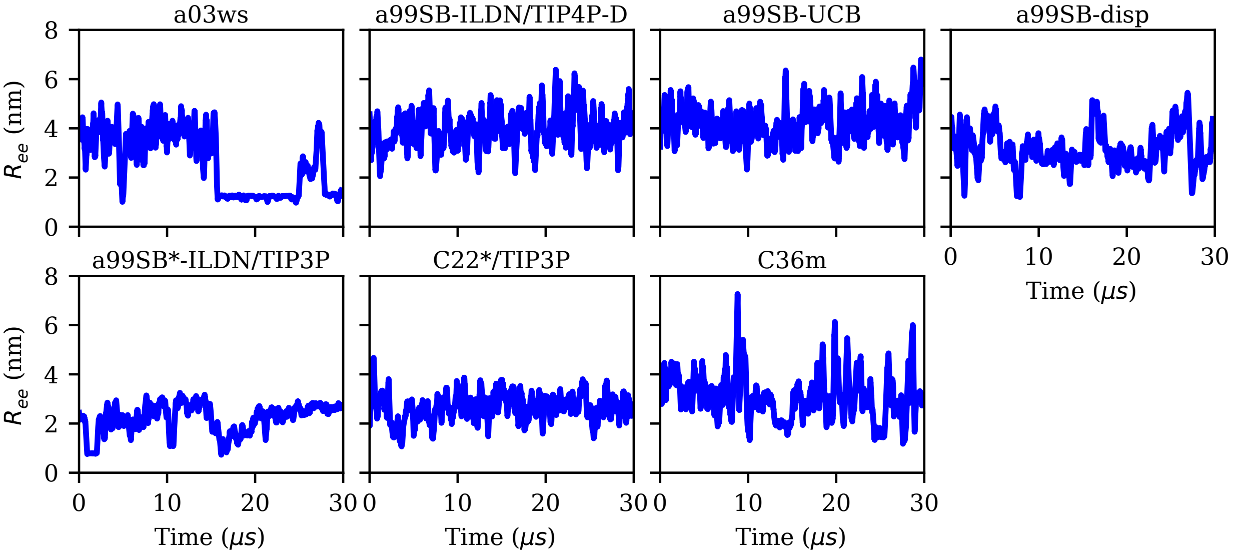


Figure S13: Evolution the end-to-end distance $R_{\mathrm{ee}}$ for the different force fields (labels on the top of the panels).

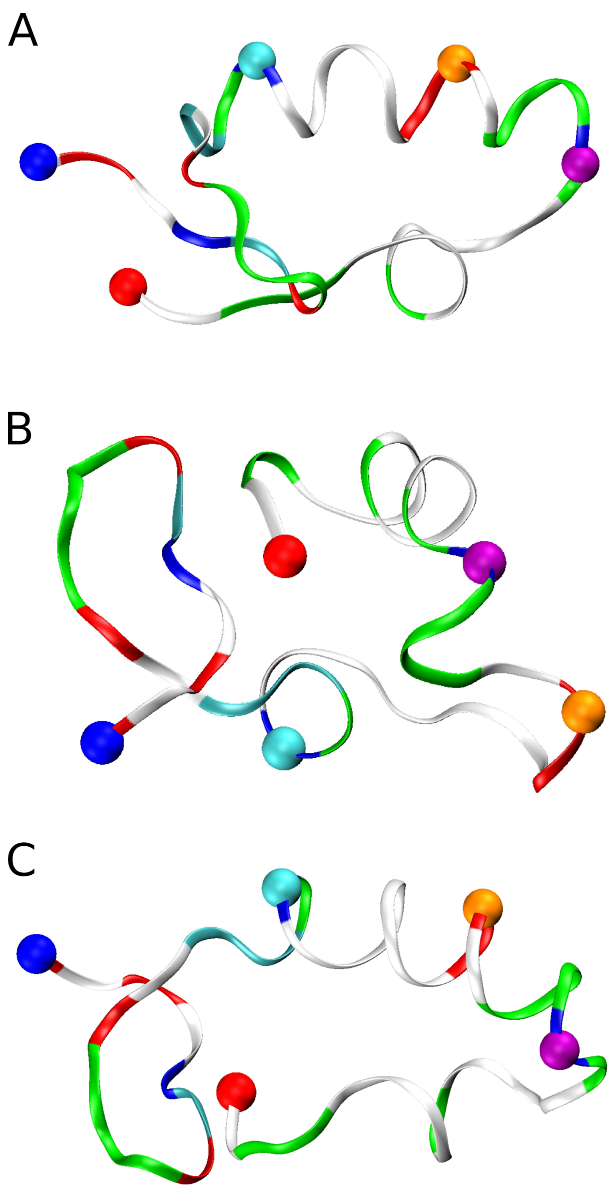


Figure S14: Compact Aβ40 structures sampled with a03ws between 16 and 25 μs. These conformations exhibit a high propenisty for helix formation in different parts along the sequence: (A) between residues K16 and K28 (as present in MSM state 1), (B) between residues G29 and M35 (as present in MSM states 3), (C) between residues K16 to K28 and G29 to M35 (as present in MSM state 2). Aβ40 is shown as band and colored according to amino acid residue type (basic: blue, acidic: red, histidine: cyan, polar: green, hydrophobic: white). Following residues are indicated by spheres: N-terminus (blue), K16 (cyan), D23 (orange), K28 (mauve), C-terminus (red).

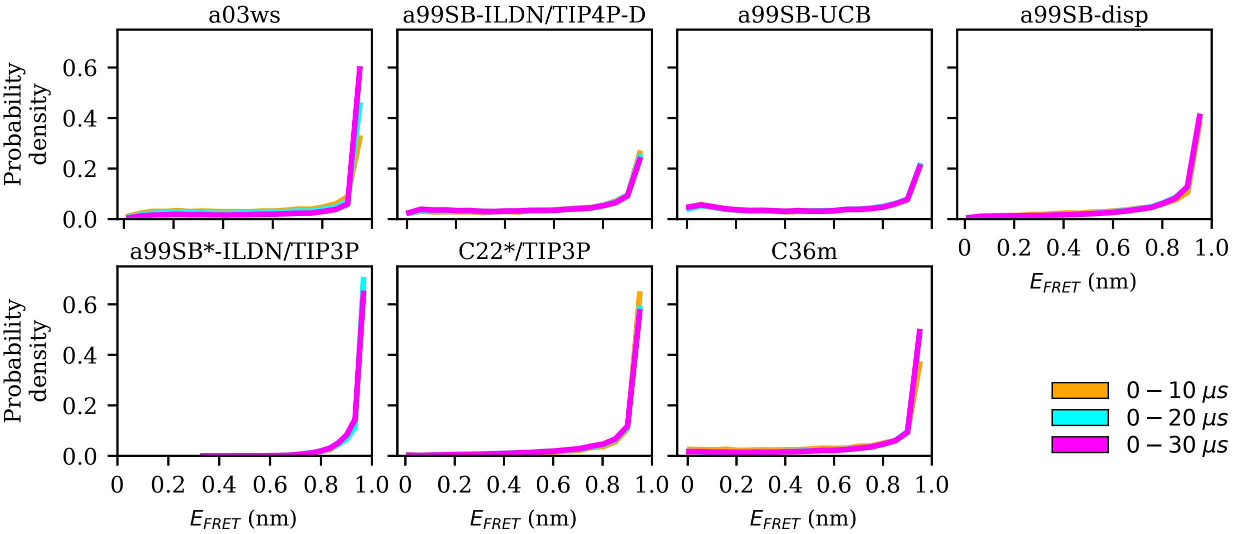


Figure S15: Distribution of the FRET efficiency $E_{\mathrm{FRET}}$ for increasing trajectory lengths (0–10 μs: yellow, 0–20 μs: cyan, 0–30 μs: magenta) for the different force fields (labels on the top of the panels).

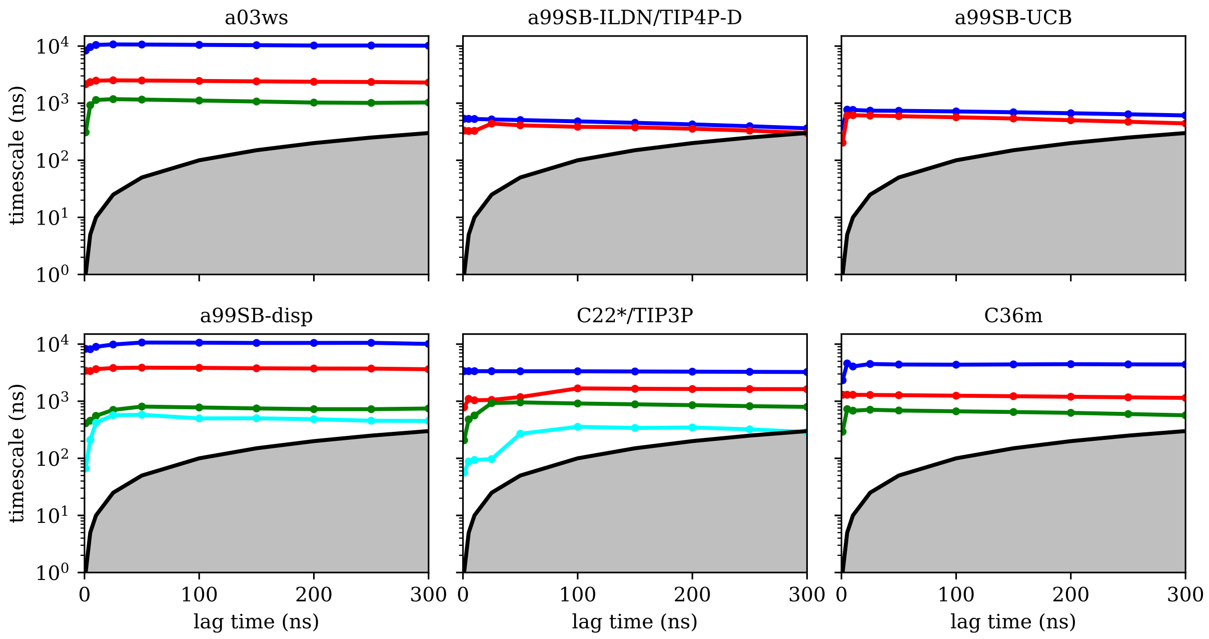


Figure S16: Implied time scales of the slowest processes (colored lines) obtained for different MSMs at different lag times (dots on colored lines) calculated from the MD trajectories using different force fields (labels on the top of the panels).

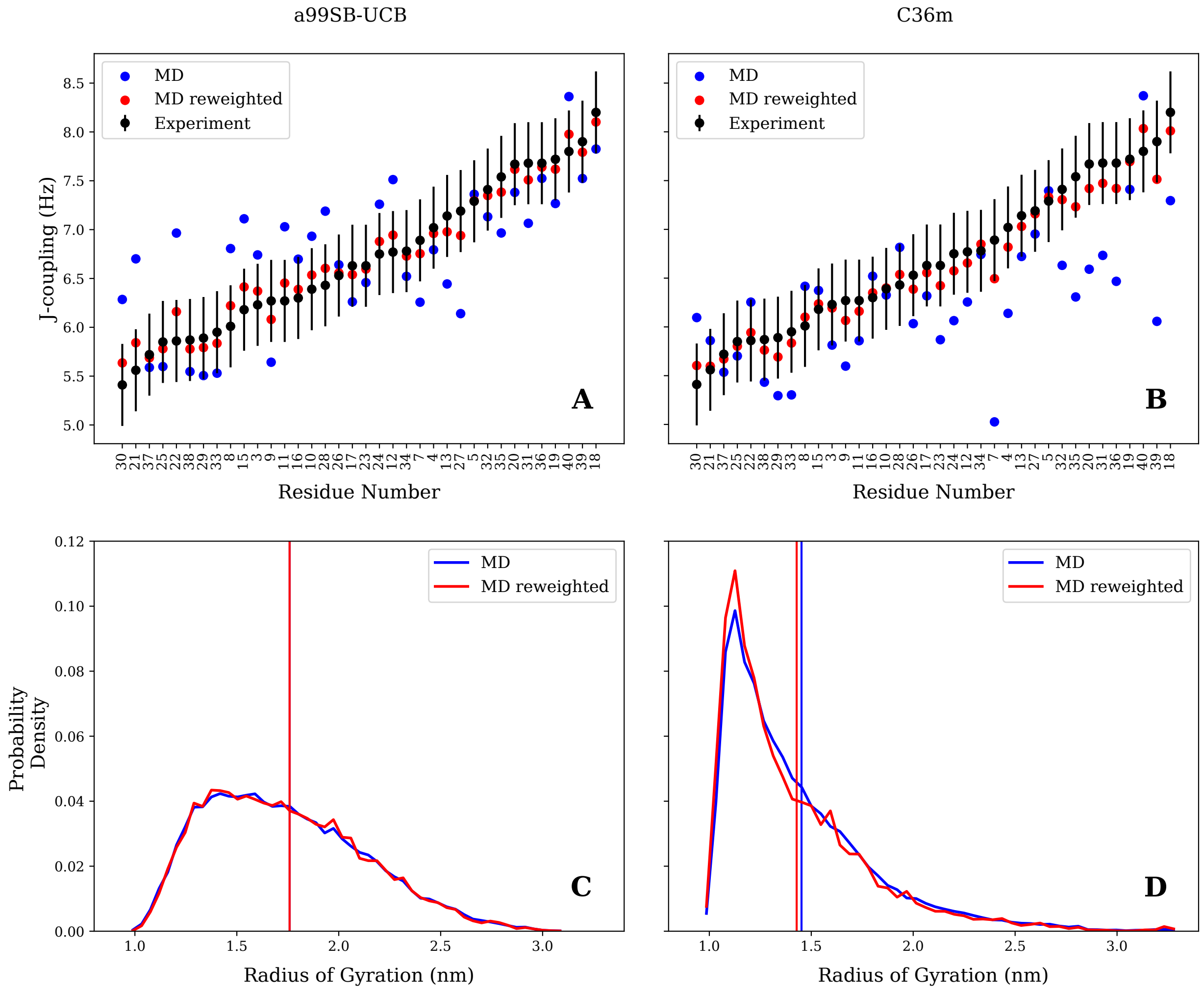


Figure S17: Reweighting of the trajectory frames using the maximum entropy principle to optimize the $J$-couplings obtained with the MD trajectory with a99SB-UCB (left) and C36m (right). (Top) The black dots indicate the experimental $J$-couplings for the individual Aβ40 residues (sorted in increasing $J$-coupling order), blue and red dots indicate the calculated $J$-couplings before and after, respectively, reweighting. (Bottom) Distribution of the radius of gyration before (blue) and after (red) reweighting the MD frames. The vertical lines indicate the corresponding $R_{\text{gyr}}$ average.

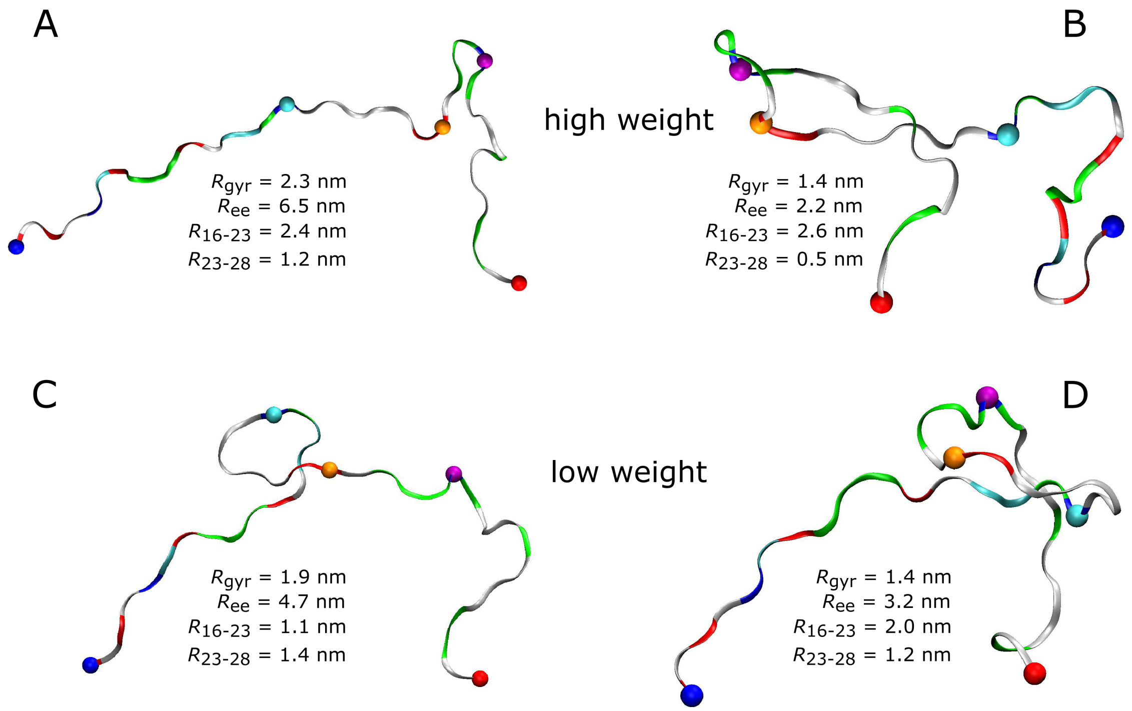


Figure S18: High-weight (A and B) and low-weight structures (C and D) determined by reweighting the C36m trajectory using the Bayesian/maximum entropy technique. Aβ40 is shown as band and colored according to amino acid residue type (basic: blue, acidic: red, histidine: cyan, polar: green, hydrophobic: white). Following residues are indicated by spheres: N-terminus (blue), K16 (cyan), D23 (orange), K28 (mauve), C-terminus (red). The structures were characterized in terms of $R_{\mathrm{ee}}$, $R_{\mathrm{gyr}}$, the K16–D23 distance ($R_{16-23}$), and the D23–K28 distance ($R_{23-28}$).
